# Comprehensive Mapping of *SGCA* Variant Effects Reveals Domain-Specific Constraints Relevant to Sarcoglycanopathies

**DOI:** 10.64898/2026.07.22.740214

**Authors:** Shushu Huang, Kenneth Ng, Yanyu Lu, Zhiying Xie, Chaoping Hu, Bochen Zhu, Wenhua Zhu, Angela Lek, Yujiao Yang, Semin Kim, Kaiyue Ma, Monkol Lek

**Affiliations:** Department of Genetics, Yale School of Medicine, New Haven, CT 06520, USA; Department of Neurology, Peking University First Hospital, Beijing, 100034, China; Department of Neurology, National Children’s Medical Center, Children’s Hospital of Fudan University, Shanghai, 201102, China; Department of Neurology, Huashan Rare Disease Center, Huashan Hospital, Fudan University, National Center for Neurological Disorders, Shanghai, 200040, China; Muscular Dystrophy Association, Chicago, IL 60607, USA; Bio-X Institutes, Key Laboratory for the Genetics of Developmental and Neuropsychiatric Disorders, Ministry of Education, Shanghai Jiao Tong University, Shanghai, 200000, China

**Keywords:** SGCA, sarcoglycanopathies, limb-girdle muscular dystrophy, variant interpretation, saturation mutagenesis, functional genomics, membrane protein

## Abstract

Pathogenic variants in *SGCA*, encoding α-sarcoglycan, cause an autosomal recessive limb-girdle muscular dystrophy, LGMDR3/2D, yet clinical interpretation of *SGCA* variants remains challenging due to the high prevalence of rare missense variants. α-sarcoglycan is an essential component of the sarcoglycan complex at the muscle cell membrane, and pathogenic variants frequently impair its membrane localization. Here, we systematically assess the effects of all possible single-nucleotide variants across the *SGCA* coding sequence using a saturation mutagenesis-based experimental assay that quantifies α-sarcoglycan surface expression. We generate a comprehensive functional atlas that distinguishes tolerated and damaging variants, aligning with independent genetic and clinical evidence, and reveals domain-specific properties of the cytoplasmic region, in which C-terminal truncating variants retain membrane localization, suggesting possible pathogenic mechanisms beyond impaired trafficking. This work provides a scalable functional framework to support genetic diagnosis and variant interpretation in sarcoglycanopathies.

**Graphical Abstract:** Schematic overview of saturation mutagenesis-based functional mapping of *SGCA*.

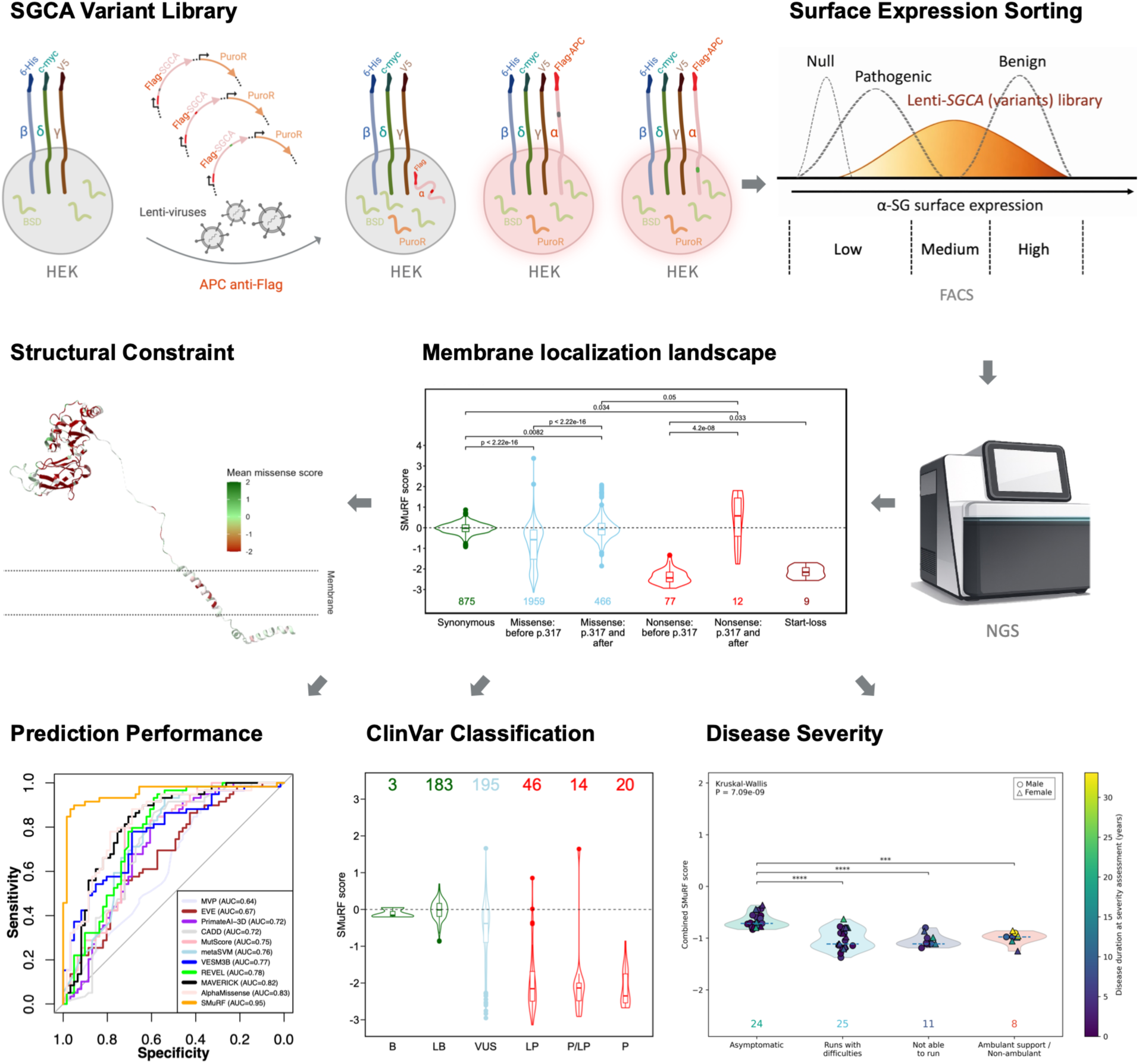

## Introduction

Sarcoglycanopathies are a group of severe, early-onset autosomal recessive limb-girdle muscular dystrophies characterized by progressive skeletal muscle weakness, loss of ambulation, and cardiopulmonary involvement. ^1,2^ These disorders arise from pathogenic variants in the sarcoglycan genes (*SGCA*, *SGCB*, *SGCD*, and *SGCG*), which encode α-, β-, δ-, and γ-sarcoglycan, respectively. Together, these proteins assemble into the sarcoglycan (SG) complex, a critical substructure of the dystrophin-glycoprotein complex (DGC) that maintains sarcolemmal stability during muscle contraction. Disruption of SG complex integrity leads to secondary destabilization of the DGC, rendering muscle fibers susceptible to contraction-induced damage.^3–6^

Among sarcoglycan genes, *SGCA* is the most frequently implicated in disease, accounting for approximately 40-60% of genetically confirmed sarcoglycanopathy cases across multiple cohorts, and exhibits a broad mutational spectrum dominated by rare missense variants.^7–11^ While next-generation sequencing (NGS) has greatly expanded the detection of *SGCA* variants in affected individuals, accurate clinical interpretation remains a major bottleneck. Nearly half (48.0%) of *SGCA* variants with ClinVar^12^ assertions are classified as variants of uncertain significance (VUS) or have conflicting interpretations (Figure S1A), limiting diagnostic certainty, genetic counseling, and eligibility for emerging gene- and variant-directed therapies.^13,14^ Despite rapid advances in artificial intelligence-driven variant interpretation, computational predictions alone are often insufficient to support clinical decision-making.^15,16^ This limitation is especially evident for membrane-associated structural proteins such as sarcoglycans, whose functions depend on complex processes including protein folding, intracellular trafficking, and SG complex assembly that may not be fully captured by current in silico models.^17^

Experimental functional assays provide a powerful and independent line of evidence for variant interpretation, particularly when deployed at scale. Deep mutational scanning approaches have demonstrated the feasibility of generating comprehensive variant effect maps; however, their widespread adoption has been constrained by technical complexity, cost, and assay design limitations.^18^ To address these challenges, we previously developed saturation mutagenesis-reinforced functional assays (SMuRF), a scalable and accessible framework that couples pooled saturation mutagenesis with quantitative, disease-relevant functional readouts to enable systematic interrogation of all possible coding variants in disease-related genes.^19^ Building on this framework, we extend saturation mutagenesis-based functional mapping to sarcoglycan genes, starting with *SGCA*.

Correct cell surface localization of the SG complex, including α-, β-, γ-, and δ-sarcoglycan, is a defining feature of sarcolemmal integrity and a hallmark routinely assessed in patient muscle biopsies.^17,20–23^ Loss of α-sarcoglycan from the cell surface is a characteristic pathological feature of SGCA-related disease; therefore, cell-surface localization represents a biologically relevant functional readout for assessing the impact of *SGCA* variants.^17,24,25^ Recent saturation mutagenesis of β-sarcoglycan (*SGCB*) demonstrated that membrane localization scores accurately predict variant pathogenicity, correlate with clinical disease severity, and provide structural insights into the SG complex assembly.^26^ The findings demonstrated cell-surface localization as a powerful functional readout for sarcoglycan variant interpretation. However, whether similar principles apply to α-sarcoglycan (*SGCA*) remains unclear. Notably, α-sarcoglycan is structurally distinct from the other sarcoglycan proteins, possessing an N-terminal signal peptide and an inverted membrane topology with an extracellular N-terminus and intracellular C-terminus,^27–29^ suggesting potential differences in protein biogenesis and variant tolerance.

In this study, we generated a comprehensive functional map of the effects of all possible single-nucleotide variants (SNVs) on α-sarcoglycan surface expression. Beyond enabling high-confidence classification, our dataset reveals unexpected domain-specific behaviors, particularly within the cytoplasmic C-terminal region. Collectively, this work establishes a comprehensive functional resource for *SGCA*, extends the application of SMuRF to membrane structural proteins, and provides a foundation for improved diagnosis and mechanistic understanding of sarcoglycanopathies.

## Results

### Construction of a Saturation Mutagenesis Library Covering All *SGCA* Single-Nucleotide Variants

To systematically characterize the functional impact of all possible coding variants in *SGCA*, we generated a saturation mutagenesis library encompassing every SNV across the full-length coding sequence (CDS). Using a pooled oligonucleotide-based mutagenesis strategy, each nucleotide position was targeted to introduce all possible SNVs, corresponding to synonymous, missense, nonsense, and start-loss substitutions (Figure S1B). Variant libraries were then cloned into a lentiviral expression vector, enabling pooled delivery and functional assessment. Adequate expression is required for robust reconstitution of the SG complex and reliable quantification of cell surface localization. Therefore, the EF1α core promoter was selected to achieve sufficient but relatively moderate expression of α-sarcoglycan, reducing the risk of overexpression-associated masking of deleterious variant effects. To enable quantitative assessment of α-sarcoglycan surface localization, an extracellular FLAG epitope tag separated by a short linker was inserted immediately downstream of the signal peptide cleavage site, preserving native signal peptide processing while allowing live-cell detection of surface-exposed protein. To facilitate short-read NGS, the *SGCA* CDS was divided into 5 non-overlapping mutagenesis blocks, each subjected to independent library construction, functional screening, and downstream analysis. A puromycin resistance cassette driven by the hPGK promoter was included to enrich for cells carrying integrated *SGCA* variant constructs prior to functional screening (Figure 1A). Variant libraries were then packaged into lentiviral particles. Deep sequencing of the plasmid pool and post-transduction genomic DNA libraries confirmed a complete representation of all possible coding SNVs (Figures S1C, S1D, and S2E), enabling high-confidence downstream analyses (Figure 2A).

**Figure 1.**
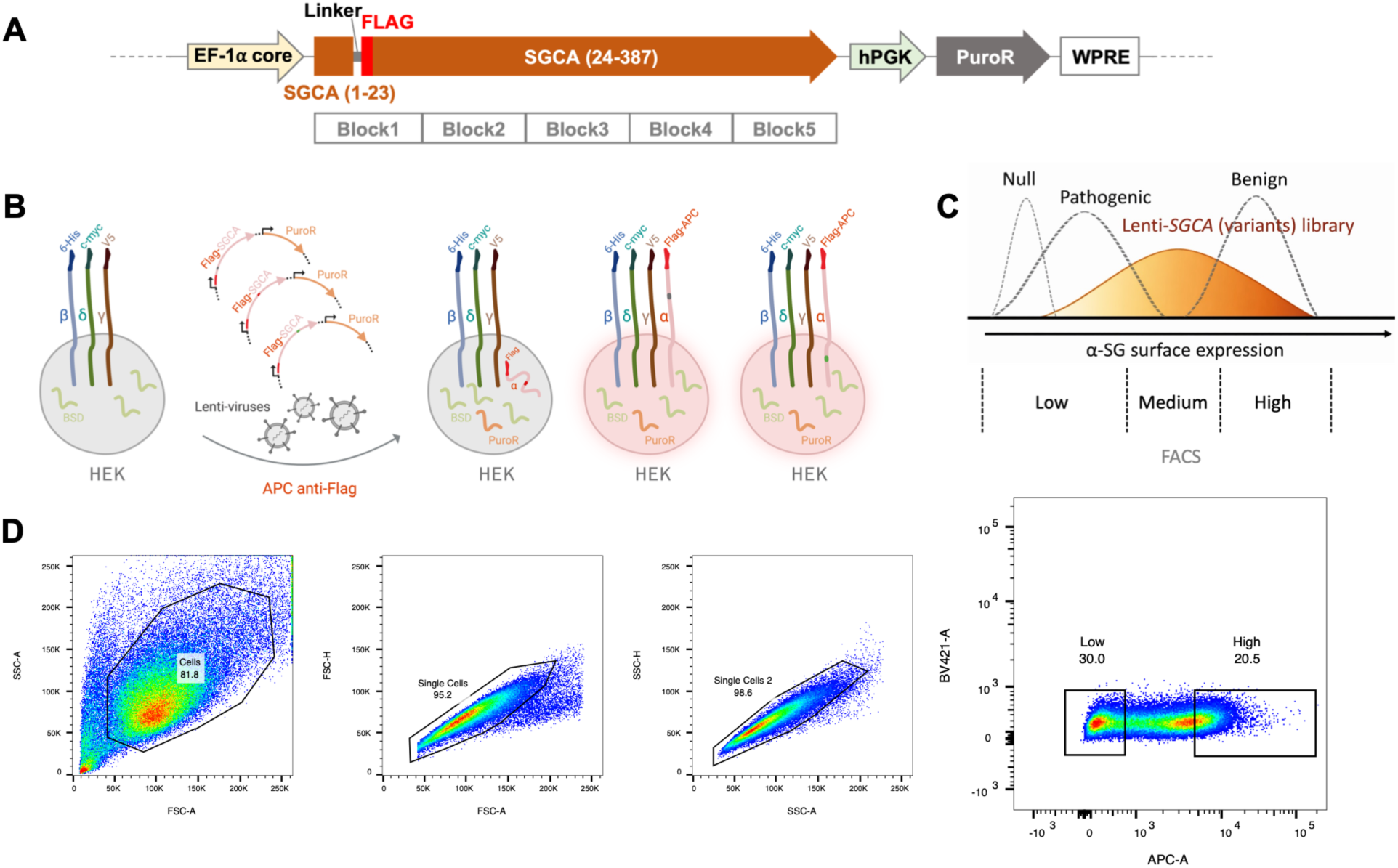
SMuRF workflow for systematic functional assessment of *SGCA* variants. **(A)** Schematic of the lentiviral expression construct, EF1A-FLAG-SGCA-hPGK-PuroR, used for saturation mutagenesis. A linker-FLAG epitope was inserted between residues 23 and 24 of SGCA, immediately downstream of the signal peptide. The coding sequence was divided into five mutagenesis blocks for library construction and sequencing. **(B)** HEK-BDG cells were transduced with lentiviral *SGCA* variant libraries, selected with puromycin, stained with APC-conjugated anti-FLAG antibody, and subjected to fluorescence-activated cell sorting (FACS). **(C)** Schematic of FACS-based separation of cells according to SGCA surface expression. **(D)** Representative FACS gating strategy used for pooled screening. Live single cells were sequentially gated and sorted into low- and high-surface-expression populations for downstream sequencing.

**Figure 2.**
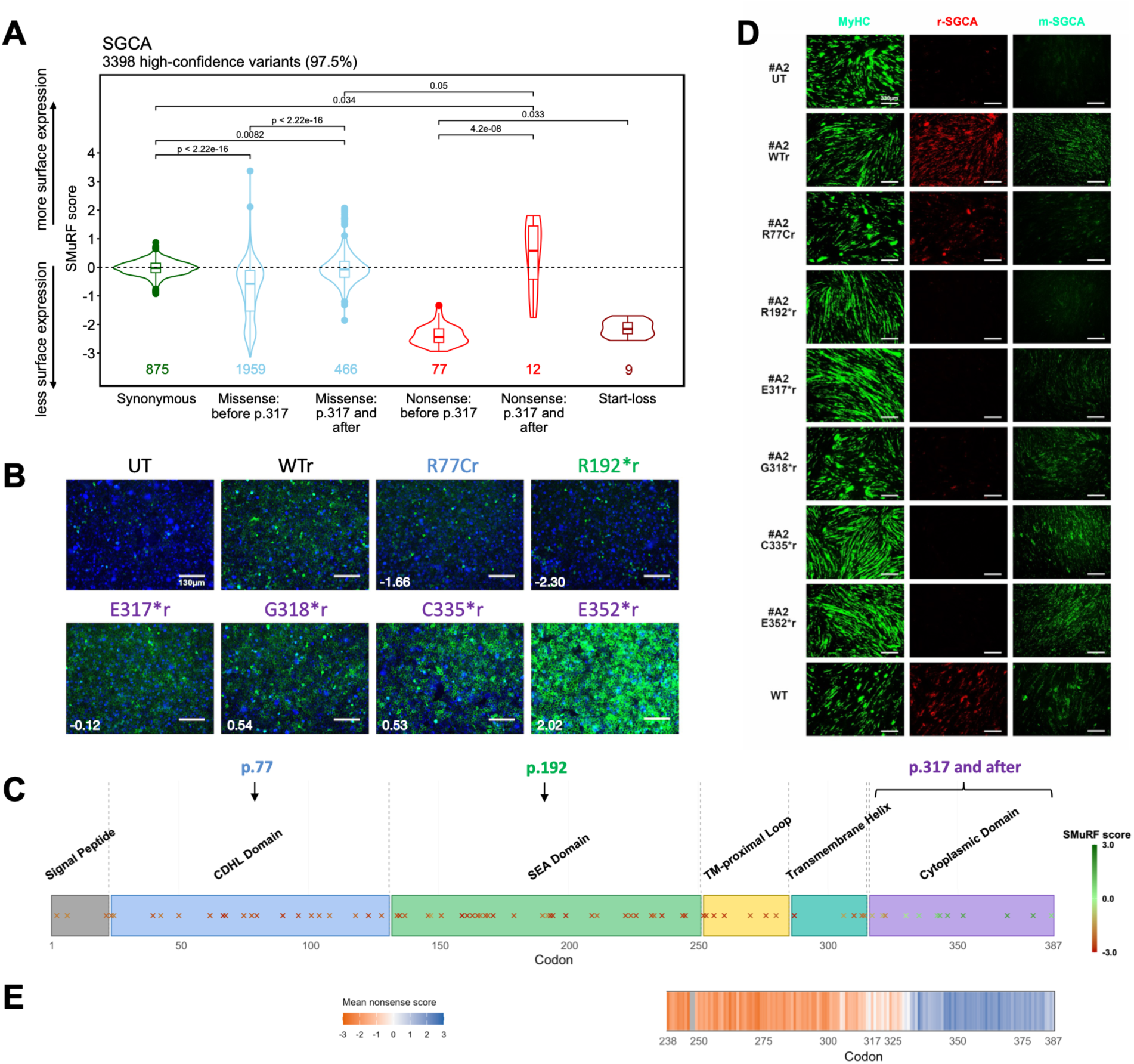
Membrane localization landscape of *SGCA* variants reveals distinct properties of the cytoplasmic domain. **(A)** High-confidence SMuRF scores align with *SGCA* variant classes. High-confidence *SGCA* variants were grouped by variant class, with missense and nonsense variants further stratified according to their position relative to residue 317. The mean of synonymous variants was used to normalize SMuRF scores. The box boundaries represent the 25th and 75th percentiles, with the horizontal line indicating the median and whiskers extending to 1.5 × the interquartile range (IQR). Variant counts are shown below each violin. P values were calculated using two-sided Wilcoxon tests. **(B)** Representative immunofluorescence images of selected *SGCA* variants. Individual *SGCA* variant constructs were transduced into HEK-BDG#9 cells using lentivirus, selected with puromycin, and analyzed by immunofluorescence. Pathogenic variants R77C and R192* served as negative controls. DAPI, blue; FLAG-SGCA, green. Numbers in the lower-left corner indicate the corresponding membrane localization scores (MLSs). Scale bars, 130 μm. **(C)** SMuRF scores of nonsense variants were mapped onto the protein domain architecture of SGCA. Scores are visualized using a color scale, with red indicating reduced membrane localization and green indicating preserved membrane localization. CDHL, cadherin-like; SEA, sea urchin sperm protein, enterokinase, agrin; TM, transmembrane. **(D)** Representative immunofluorescence images of MB135 wild-type (WT) myotubes or MB135 *SGCA*-knockout (KO) (clone A2) myotubes expressing WT or indicated *SGCA* variants. Fixed myotubes were stained with MyHC and two independent SGCA antibodies recognizing distinct regions of the protein: r-SGCA, directed against the transmembrane/cytoplasmic region (aa288-387), and m-SGCA, directed against the extracellular domain (aa217-289). Scale bars, 330 μm. **(E)** Targeted nonsense-only saturation mutagenesis of the C-terminal region of *SGCA*. A targeted nonsense library was generated by introducing a premature stop codon TAA at every codon between residues 238 and 387. The heatmap displays the mean score for nonsense variants from two biological replicates (lentiviral transduction in HEK-BDG #425 and #426) at each codon, with orange indicating reduced membrane localization and blue indicating preserved membrane localization. Membrane localization increased progressively across a transition zone spanning approximately residues 315-333, rather than exhibiting an abrupt boundary at residue 317. Residue 317 is indicated on the x-axis. Positions with low-confidence mean nonsense scores are shown in gray. Low-confidence positions were defined as those with fewer than 100 total reads (high + low populations) in at least one replicate or an inter-replicate standard deviation > 1.5. The heatmap is presented on the same amino acid scale as the SGCA domain architecture in panel C to facilitate visualization of domain-specific effects.

### Quantitative Measurement of α-Sarcoglycan Surface Expression Using a Cell-Based FACS Assay

Functional consequences of *SGCA* variants were assessed using a cell-based assay that directly reports α-sarcoglycan surface expression. HEK293T cells, which lack endogenous expression of sarcoglycan proteins, have been widely used in previous studies of SG complex and their variant effects, supporting their suitability as a platform for establishing a controlled sarcoglycan expression system.^26,30–32^ To recapitulate SG complex formation, we generated an engineered HEK293T-derived cell line (HEK-BDG) stably expressing β-, δ-, and γ-sarcoglycan. Lentiviral delivery of the pooled *SGCA* variant libraries into this background enabled systematic assessment of variant effects with incorporation of the SG complex (Figure 1B).

Live-cell staining against the extracellular epitope FLAG tag was performed under non-permeabilizing conditions, followed by fluorescence-activated cell sorting (FACS) to separate populations with High, Medium, and Low surface expression. Sorted populations with High and Low surface expression were collected at a minimum coverage of ~1,500× to maintain robust variant representation (Figures 1C and 1D). Genomic DNA was extracted from each population, and variant frequencies were quantified by NGS. A membrane localization score (MLS) was calculated as the normalized log-enrichment of each variant between high- and low-expression populations relative to the wild-type sequence, yielding a continuous quantitative measure of surface localization capacity.

### SMuRF Score Distributions Reveal Position-Dependent Effects Across *SGCA* Variant Classes

Three biological transduction replicates were performed using three independent clonal HEK-BDG stable cell lines to confirm the reproducibility (Figures S2A-C). Pairwise comparison of replicate-specific log2 MLS demonstrated strong concordance across biological replicates (Pearson r = 0.75-0.81) (Figure S2D). DiMSum^33^ was used to combine results from three biological replicates, and SMuRF scores, representing the combined MLSs, were generated for all possible coding SNVs of *SGCA* (3,483) (Figure S2E), of which 3,398 (97.6%) met the high-confidence threshold and were retained for downstream analyses (Figure 2A).

To investigate the effects of different classes of *SGCA* variants on membrane localization, we analyzed the high-confidence variants and stratified them by molecular consequence and protein position (Figure 2A). Synonymous variants (n = 875) clustered near wild-type levels, with a median SMuRF score of −0.022 (95% CI: −0.036 to −0.001) and a central 95% distribution range of −0.593 to 0.476, establishing the reference distribution for variants with negligible functional impact. Conversely, start-loss variants (n = 9) were strongly deleterious, with a median SMuRF score of −2.158 (95% CI: −2.449 to −1.834) and a central 95% distribution range of −2.539 to −1.724. Missense variants demonstrated a pronounced position-dependent effect. Variants occurring before p.317 (n = 1,959) showed a lower median SMuRF score (−0.578; 95% CI: −0.640 to −0.532) and a broad central 95% distribution range (−2.704 to 0.698), spanning nearly the full dynamic range of the assay and enabling resolution between tolerated and damaging variants. In contrast, missense variants at and after p.317 (n = 466) were shifted toward wild-type-like membrane localization, with a median score of −0.075 (95% CI: −0.124 to −0.028) and a central 95% distribution range of −0.981 to 1.073. As p.317 represents the first assayed residue within the SGCA cytoplasmic domain (residue 316-387), these findings suggest that amino acid substitutions within the cytoplasmic tail are generally better tolerated with respect to membrane localization than those affecting the extracellular and transmembrane regions. Lastly, nonsense variants exhibited an even more pronounced positional effect. Nonsense variants before p.317 (n = 77) were strongly deleterious, with a median SMuRF score of −2.437 (95% CI: −2.535 to −2.317) and a central 95% distribution range of −2.878 to −1.567. In contrast, nonsense variants at and after p.317 (n = 12) showed substantially higher scores, with a median of 0.577 (95% CI: −0.586 to 1.510) and a central 95% distribution range of −1.556 to 1.775, indicating that many C-terminal truncating variants retain near-wild-type membrane localization. Together, these results indicate that the extracellular and transmembrane regions of SGCA are considerably less tolerant to variation than the cytoplasmic domain, particularly with respect to truncating variants.

### C-Terminal Truncating Variants Retain Surface Expression Despite Loss of the Cytoplasmic Domain

Individual validation confirmed the unexpected tolerance of C-terminal truncating variants identified in the pooled screen. Representative nonsense variants located at or beyond p.317, together with wild-type SGCA and two pathogenic controls (R77C and R192*), were individually expressed in HEK-BDG cells and evaluated by live-cell staining of the extracellular FLAG epitope. Consistent with the MLSs obtained from pooled screening, C-terminal truncating variants retained robust cell-surface localization, with staining patterns comparable to or exceeding those of wild-type SGCA. In contrast, R77C and the early truncating variant R192* exhibited markedly reduced surface expression (Figures 2B and 2C).

To determine whether this phenotype extended beyond the engineered HEK-BDG platform, we generated *SGCA*-knockout (KO) MB135 human myoblasts (Figures S3A-C) and reintroduced individual *SGCA* variants prior to differentiation into myotubes as an orthogonal validation. Staining with an antibody recognizing the extracellular domain of SGCA (m-SGCA) closely recapitulated both the HEK-BDG localization phenotypes and the MLSs obtained from pooled screening. C-terminal truncating variants retained robust extracellular-domain staining, whereas R77C and R192* displayed substantially reduced signal (Figure 2D), confirming that these variants retain membrane localization in both engineered and myogenic cellular contexts. In contrast, staining with the transmembrane/cytoplasmic antibody (r-SGCA) was readily detected only in wild-type-rescued cells and was largely absent in all truncating variants (Figure 2D). This observation is consistent with loss of the corresponding antibody epitope and/or structural alterations that prevent antibody recognition of the truncated proteins. A low level of residual staining was observed for R77C, in agreement with previous reports that this common founder variant retains partial α-sarcoglycan function despite impaired trafficking and reduced protein stability.^7,11,30^

To assess whether this observation reflected a general property of the cytoplasmic domain rather than a few representative variants, we generated a nonsense variant library to introduce premature stop codons (TAA) to each codon spanning the last 150 amino acids of SGCA and examined the resulting MLSs. The nonsense library includes the distal SEA (<u>s</u>ea urchin sperm protein, <u>e</u>nterokinase, <u>a</u>grin) domain, the transmembrane (TM)-proximal Loop, the TM helix, and the cytoplasmic domain (Figure 2E). Premature stop codons within the extracellular region represented in the nonsense library markedly reduced membrane localization, whereas truncating variants within the cytoplasmic domain consistently retained wild-type-like MLSs. Rather than exhibiting an abrupt boundary at residue 317, membrane localization increased progressively across a transition zone spanning approximately residues 315-333 immediately distal to the TM helix, which is consistent with our observation of nonsense variants within the SNV library (Figure 2C). The nonsense library results indicate that preservation of membrane localization is a property of cytoplasmic truncating variants.

Together, loss of the C-terminal cytoplasmic region does not necessarily impair α-sarcoglycan’s membrane localization. Two possible interpretations can be raised. First, some C-terminal truncating variants may be less deleterious than predicted based solely on their truncating nature. Alternatively, pathogenicity may result from defects not captured by the membrane localization assay. Potential mechanisms include disruption of intracellular protein-protein interactions (e.g., γ-/δ-SG^34,35^), altered post-translational regulation (e.g., α-/γ-SG^36^), or impaired signaling functions (e.g., γ-SG^37,38^) mediated by the cytoplasmic domain.

### SMuRF Scores Align with ClinVar and Population Variation and Outperform Computational Predictors

To evaluate the clinical relevance of the *SGCA* SMuRF scores, we first examined their relationship with ClinVar^12^ classifications. Variants were grouped according to ClinVar assertions, including benign (B), likely benign (LB), variants of uncertain significance (VUS), likely pathogenic (LP), pathogenic/likely pathogenic (P/LP), and pathogenic (P). SMuRF scores were strongly concordant with clinical classification, with B and LB variants clustering near wild-type levels, whereas P, P/LP, and LP variants exhibited substantially reduced scores (Figure 3A). As expected, VUS displayed a broad distribution spanning both benign-like and pathogenic-like membrane localization profiles. Excluding variants at and beyond residue 317 resulted in even stronger concordance between SMuRF scores and ClinVar classifications (Figure S4A).

**Figure 3.**
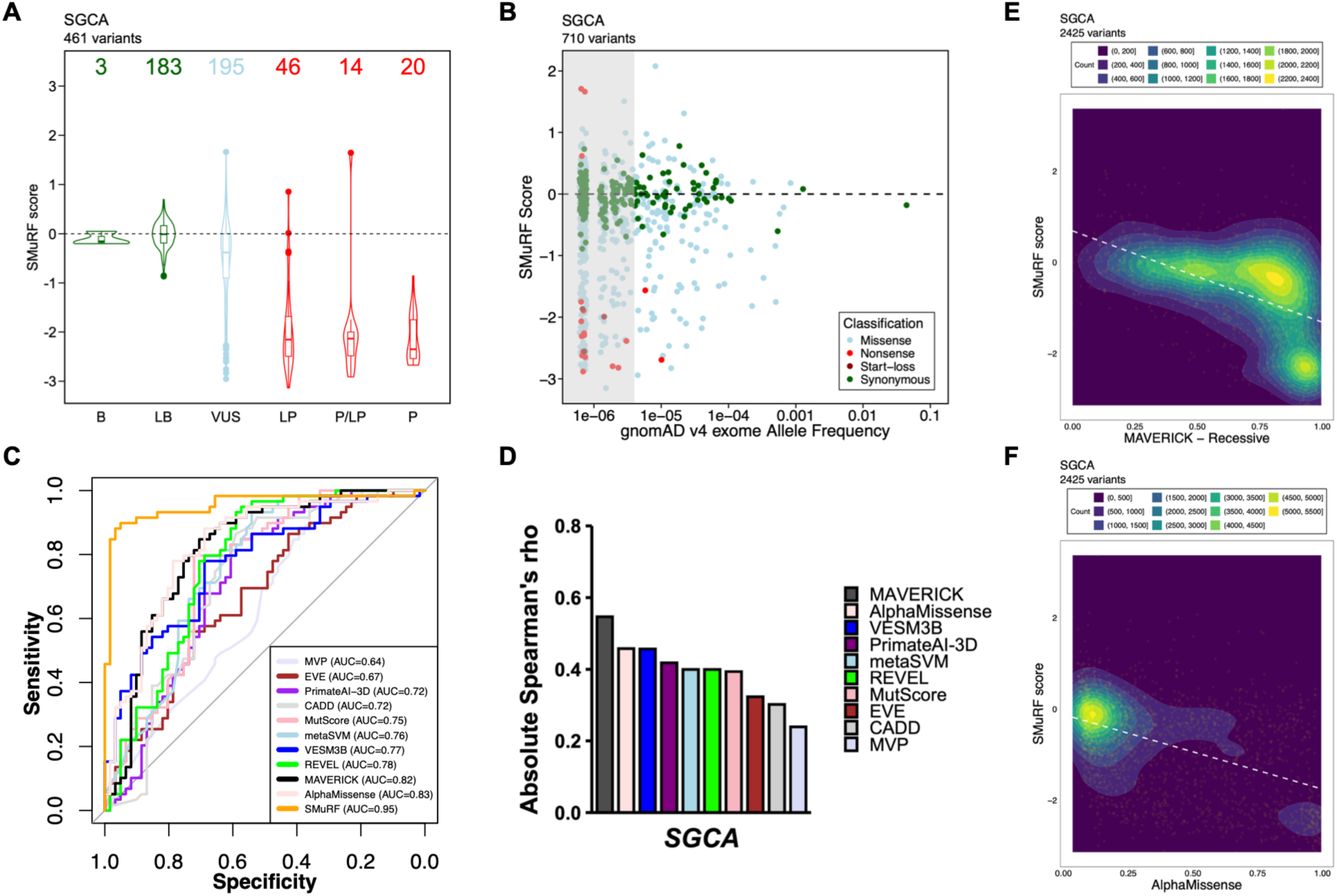
SMuRF scores align with ClinVar and population variation and outperform computational predictors. **(A)** SMuRF scores correlate well with ClinVar classifications. B, benign; LB, likely benign; VUS, variant of uncertain significance; LP, likely pathogenic; P/LP, pathogenic/likely pathogenic; P, pathogenic. Variant counts are shown above each violin. The box boundaries represent the 25th and 75th percentiles, with the horizontal line indicating the median and whiskers extending to 1.5 × the interquartile range (IQR). **(B)** SMuRF scores correlate with allele frequencies in gnomAD v4 exomes. Low-frequency variants exhibited a broad range of SMuRF scores, whereas high-frequency variants converged toward wild-type-like scores, consistent with purifying selection (Gray box, AF < 4e-6). Variants are colored according to variant class. The dashed line indicates the wild-type SMuRF score. Points were jittered for visualization. **(C)** Receiver operating characteristic (ROC) curves of SMuRF and computational predictors. The P, P/LP, and LP missense variants in ClinVar were used as the positive reference set; the negative reference set used missense variants observed in the gnomAD v4 exomes sequencing database with an AF > 1e-5, excluding any variants annotated as P, P/LP, or LP in ClinVar. AUC, area under the curve. **(D)** Absolute Spearman correlation coefficients between SMuRF scores and computational predictor scores across high-confidence *SGCA* missense variants. **(E, F)** Density plot showing the relationship between MAVERICK recessive (E) or AlphaMissense (F) scores and SMuRF scores for high-confidence *SGCA* missense variants. Color intensity represents local variant density. White dashed lines represent linear regression.

Several notable exceptions were observed (Figure S4B). The C-terminal truncating variant E352* retained a high SMuRF score (1.64) despite its P/LP classification in ClinVar, consistent with its preserved membrane localization in our individual validation experiments (Figures 2B and 2D) and nonsense library (Figure 2E). Notably, according to ClinVar (variation ID: 550601), the current classification of E352* is largely based on its predicted truncating nature and extreme rarity in the general population (Allele Frequency = 8.482e-7 in gnomAD v4.1.1), as no individuals with SGCA-related disease carrying E352* have been reported in the literature to date; therefore, direct clinical evidence supporting its pathogenicity remains limited. We also identified an additional outlier variant, R110L, which demonstrated wild-type-like membrane localization (SMuRF score = 0.85) despite having been reported in trans with the pathogenic founder variant R77C in an affected individual.^7^ However, no additional clinical or segregation data are currently available for this case. The synonymous variant L249L, a ClinVar LP variant which has been observed in patients and is predicted to disrupt normal splicing (SpliceAI^39^ Δ = 0.65, donor loss), also exhibited a wild-type-like SMuRF score (0.01). As the assay was performed using *SGCA* CDS constructs, pathogenic effects mediated through aberrant splicing are not captured, explaining the preserved SMuRF score despite its clinical classification. Supporting this interpretation, variants predicted by SpliceAI to affect splicing (SpliceAI Δ ≥ 0.5) did not exhibit significantly different SMuRF scores from variants without predicted splice effects (SpliceAI Δ < 0.5) (Figure S4E). SpliceAI delta scores and corresponding predicted splice consequences for all SNVs are provided in the “spliceai_delta” and “spliceai_consequence” columns, respectively, in Table S1.

As an independent line of evidence, we compared SMuRF scores with population allele frequencies from the Genome Aggregation Database (gnomAD) v4 exomes database. Variants with severely reduced derived SMuRF scores were concentrated toward the low-frequency end of the allele-frequency (AF) spectrum, whereas synonymous and membrane-localization-tolerated variants were distributed across a broader range of allele frequencies (Figure 3B). Notably, many variants with severely reduced SMuRF scores clustered within the ultra-rare frequency range corresponding to variants observed in five or fewer alleles in gnomAD v4 exomes (allele count ≤ 5), consistent with purifying selection against *SGCA* variants that impair membrane localization, the key pathogenic mechanism underlying dysfunction of this membrane structural protein.

In addition to supporting clinical variant interpretation, experimentally derived SMuRF scores provide an independent benchmark for evaluating computational pathogenicity predictors. We compared the performance of several widely used prediction algorithms, including AlphaMissense,^40^ MAVERICK,^41^ REVEL,^42^ MetaSVM,^43^ VESM3B,^44^ MutScore,^45^ PrimateAI-3D,^46^ CADD,^47^ EVE,^48^ and MVP,^49^ using ClinVar P, P/LP, and LP missense variants as the positive reference set. Since only two *SGCA* missense variants are currently classified as B or LB in ClinVar, common missense variants observed in gnomAD v4 exomes (AF > 1e-5) were used as a presumed benign reference set after excluding ClinVar P, P/LP, or LP variants. This threshold corresponds to approximately AC ≥ 15 and yields similarly sized positive (n = 59) and negative (n = 61) sets. Receiver operating characteristic (ROC) analysis demonstrated that SMuRF scores outperformed all computational predictors in distinguishing pathogenic variants from the presumed benign population reference set (AUC = 0.95; Figure 3C). Among computational methods, AlphaMissense showed the strongest performance (AUC = 0.83), followed by MAVERICK (AUC = 0.82), whereas other predictors achieved more modest predictive accuracy (Figure 3C). Although computational predictors provide useful evidence for variant interpretation, these results indicate that direct experimental measurement of a disease-relevant cellular phenotype offers substantially greater predictive capability for *SGCA* variant classification.

To assess concordance between computational predictions and experimentally measured membrane localization, we calculated Spearman correlations between SMuRF scores and predictor scores across all high-confidence *SGCA* missense variants. Correlation strengths varied considerably among predictors (Figure 3D). MAVERICK exhibited the strongest correlation with SMuRF scores, followed by AlphaMissense and VESM3B, whereas other predictors showed weaker agreement with experimental measurements. MAVERICK recessive scores demonstrated an overall inverse relationship with SMuRF scores, with variants predicted to be more damaging generally exhibiting reduced membrane localization. Although showing stronger overall agreement with SMuRF, a substantial subset of variants assigned high MAVERICK scores retained near wild-type membrane localization (Figure 3E). Similarly, AlphaMissense scores showed an overall inverse relationship with SMuRF scores (Figure 3F); however, most variants received low AlphaMissense scores, yet these variants exhibited a broad range of SMuRF scores, including a subset with markedly reduced membrane localization. VESM3B scores showed an overall positive relationship with SMuRF scores, although did not fully resolve variants with distinct membrane localizations (Figure S4C). Together, these results highlight the complementary value of experimental functional measurements for resolving variant effects beyond current computational predictions.

### The *SGCA* Variant Effect Map Reveals Domain- and Structure-Specific Functional Constraints

To investigate how functional constraint is distributed across α-sarcoglycan, we calculated a residue-level mean missense score, defined as the mean SMuRF score of all missense substitutions at each amino acid position. These mean missense scores were visualized on both the linear human SGCA protein sequence (Figure 4B) and the recently resolved rabbit cryo-EM structure of the α-sarcoglycan within DGC^28^ (Figure 4A), providing a residue-level map of substitution tolerance across the SGCA protein. The independently determined mouse SGCA structure^29^ exhibited a highly similar overall architecture but, like the rabbit structure, did not fully resolve the cytoplasmic domain structure (Figure S5A). An AlphaFold^50,51^ predicted human SGCA model was therefore used to visualize the complete protein (Figure S5B).

**Figure 4.**
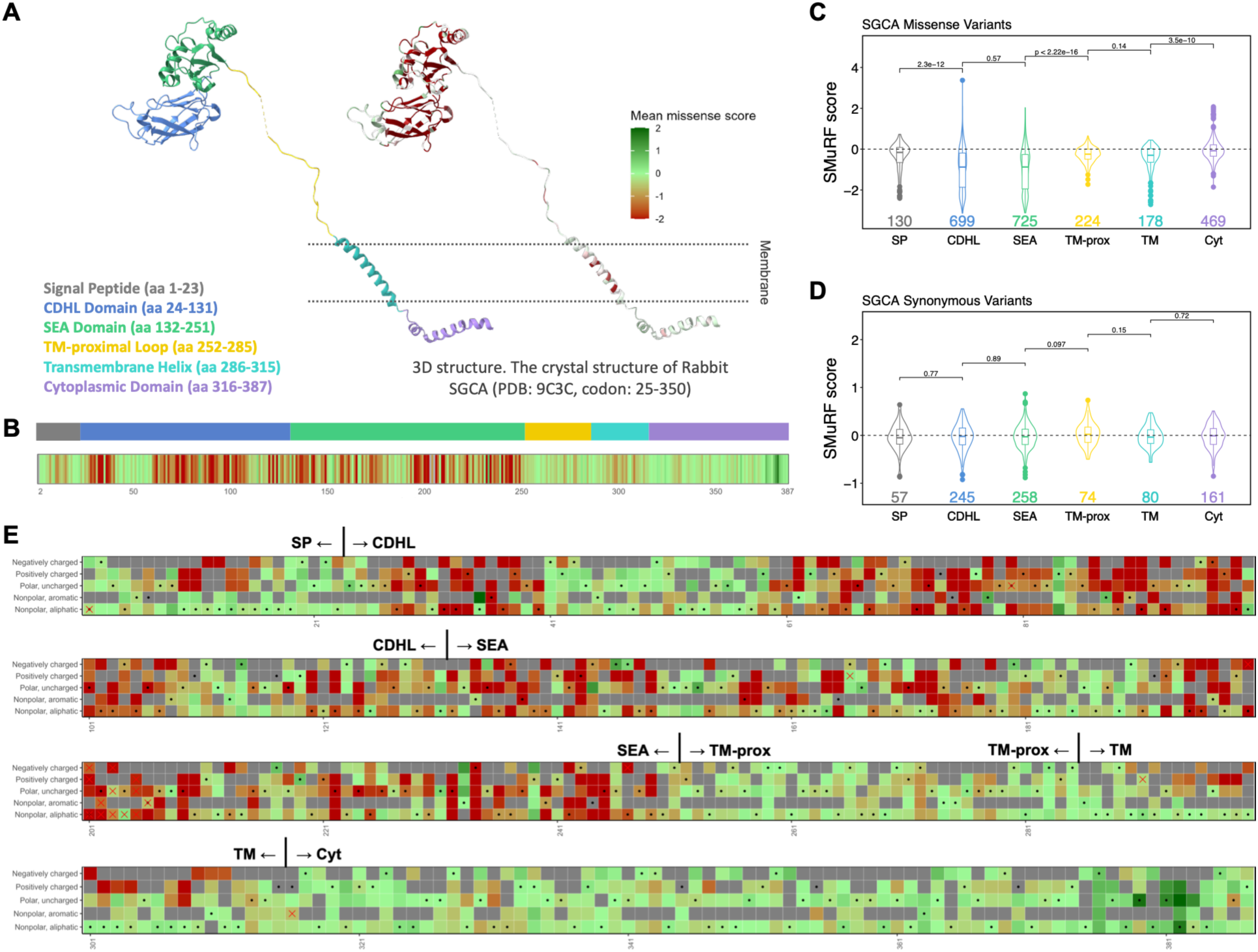
The *SGCA* Variant Effect Map Reveals Domain- and Structure-Specific Functional Constraints. **(A)** Residue-level mean missense SMuRF scores mapped onto the cryo-EM structure of the rabbit á-sarcoglycan within the dystrophin-glycoprotein complex (DGC; PDB: 9C3C). Mean missense scores were calculated as the average SMuRF score of all missense substitutions at each amino acid position. Residues are colored according to the mean missense score, with red indicating reduced membrane localization and green indicating preserved membrane localization. Structural domains are annotated according to the linear protein sequence. **(B)** Residue-level mean missense SMuRF scores mapped onto the linear SGCA protein sequence. Colors represent mean missense SMuRF scores using the same scale as in panel A. Protein domains are shown above the heatmap. **(C, D)** Distribution of SMuRF scores for missense (C) and synonymous (D) variants across SGCA protein domains. Violin plots show the distribution of SMuRF scores, with box boundaries representing the 25th and 75th percentiles, the horizontal line indicating the median, and whiskers extending to 1.5 × the interquartile range (IQR). Variant counts are shown below each violin. P values were calculated using two-sided Wilcoxon tests. **(E)** Position-specific visualization of missense variant effects grouped by amino acid physicochemical properties. Amino acid substitutions were classified into five groups: negatively charged (D, E), positively charged (K, R, H), polar uncharged (S, T, C, P, N, Q), nonpolar aromatic (F, Y, W), and nonpolar aliphatic (G, A, V, L, I, M). Each cell represents the mean SMuRF score of all single-nucleotide variant (SNV)-accessible substitutions within the corresponding amino acid class at a given codon. Colors represent mean missense SMuRF scores using the same scale as in panel A. Black dots indicate the physicochemical class of the wild-type residue. Gray squares indicate amino acid substitutions that cannot be generated by a single-nucleotide substitution. Red crosses indicate positions at which all SNV-accessible substitutions within the corresponding physicochemical class were classified as low confidence. Abbreviations: SP, signal peptide; CDHL, cadherin-like; SEA, sea urchin sperm protein, enterokinase, agrin; TM-prox, transmembrane-proximal loop; TM, transmembrane helix; Cyt, cytoplasmic domain.

Distinct patterns of functional constraint were observed across the protein. Regions corresponding to the extracellular CDHL (<u>c</u>a<u>dh</u>erin-like) and SEA domains were enriched for residues with strongly reduced mean missense scores, indicating that amino acid substitutions at these positions are generally poorly tolerated. The signal peptide and TM helix also exhibited appreciable functional constraint, whereas the TM-proximal loop was considerably more tolerant to amino acid substitution. In contrast, the cytoplasmic C-terminal domain displayed substantially higher mean missense scores, indicating markedly greater tolerance to sequence variation.

To quantitatively compare variant tolerance across domains, SMuRF score distributions were examined separately for missense and synonymous variants (Figures 4C and 4D). Consistent with the residue-level analyses, missense variants exhibited marked domain-specific differences. In contrast, synonymous variants showed relatively uniform SMuRF score distributions across domains, indicating that the domain-specific patterns observed for missense variants reflect biological constraints on α-sarcoglycan structure and function rather than technical biases associated with variant measurement.

To facilitate interpretation of individual variants, amino acid substitutions were further grouped into five physicochemical classes: negatively charged, positively charged, polar uncharged, non-polar aromatic, and non-polar aliphatic, and visualized across the SGCA protein sequence (Figure 4E). This representation provides a position-specific view of substitution tolerance and highlights regions in which diverse classes of amino acid changes are poorly tolerated. Notably, wild-type residues within both the signal peptide and TM helix were predominantly non-polar aliphatic amino acids, consistent with the strong hydrophobic constraints required for endoplasmic reticulum (ER) targeting and membrane integration. In contrast, residues within the extracellular CDHL and SEA domains frequently exhibited intolerance to substitutions across multiple physicochemical classes, indicating that variant effects in these domains cannot be explained solely by simple physicochemical requirements.

Together with the residue-level and domain-level analyses, these findings demonstrate that functional constraints differ fundamentally between the extracellular/transmembrane regions and the cytoplasmic domain of α-sarcoglycan. These observations motivated the development of a region-specific framework for interpreting *SGCA* SMuRF scores, in which variants within residues 1-315 (extracellular/transmembrane regions) and residues 316-387 (cytoplasmic domain) are interpreted separately according to their distinct membrane-localization properties. Within residues 1-315, ClinVar B, LB, together with P, P/LP, and LP variants were used as reference groups to derive a SMuRF score threshold. Kernel density estimation, ROC analysis, and minimum classification error analysis consistently supported a SMuRF score threshold of approximately −0.9. Within residues 1-315, variants with SMuRF scores > −0.9 were classified as Functional, those with scores ≤ −0.9 were classified as Damaging, while variants within residues 316-387 were classified as membrane localization-preserved (ML-preserved) or Damaging using the same score threshold, based on the concern that membrane localization alone may not fully capture the functional consequences of variants within the cytoplasmic domain. This region-specific interpretation framework was subsequently applied to all high-confidence *SGCA* SNVs for a potential variant classification (Table 1). Beyond facilitating interpretation of experimentally measured SMuRF scores, this framework provides a practical basis for incorporating functional evidence into clinical variant interpretation, particularly under the PS3/BS3 criteria according to the standards and guidelines developed jointly by the American College of Medical Genetics (ACMG) and the Association for Molecular Pathology (AMP).^52,53^

**Table 1.**
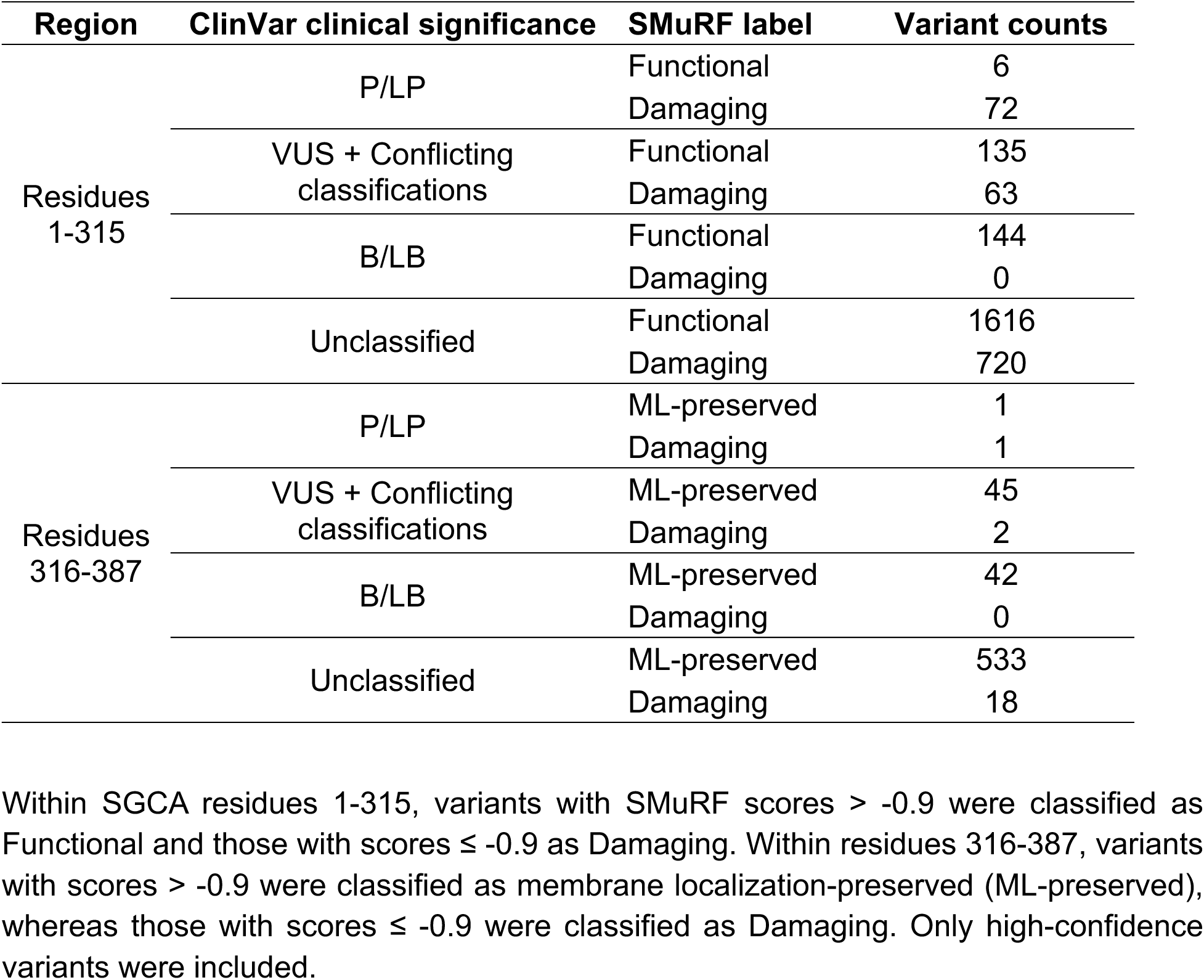
Region-specific framework for interpreting *SGCA* SMuRF scores.

### SMuRF Scores Correlate with Clinical Disease Severity and Muscle Function

To determine whether SMuRF scores capture clinically meaningful differences in disease severity, we analyzed a well-characterized clinical cohort of 68 SGCA patients assembled by collaborating neuromuscular centers in China and the USA. All individuals exhibited SGCA-associated manifestations ranging from asymptomatic hyperCKemia to severe limb-girdle muscular dystrophy and carried *SGCA* genotypes considered disease-causing by the clinical investigators. To estimate the overall effect of biallelic variants, we calculated a combined SMuRF score for each patient using both alleles as described in Methods. Patients were stratified into four disease severity categories: (1) asymptomatic hyperCKemia or minimally affected, (2) ambulatory with difficulty running, (3) ambulatory but unable to run, and (4) ambulatory with support or non-ambulant. Combined SMuRF scores progressively decreased with increasing disease severity (Figure 5A) and were strongly inversely correlated with disease severity (Spearman’s ρ = −0.67, p = 2.82e-10, N = 68; Figure S6A). Within each disease severity category, combined SMuRF scores were comparable between males and females (all BH-adjusted P > 0.5; Figure S6B). These findings support an association between reduced α-sarcoglycan membrane localization capacity and increased disease severity.

**Figure 5.**
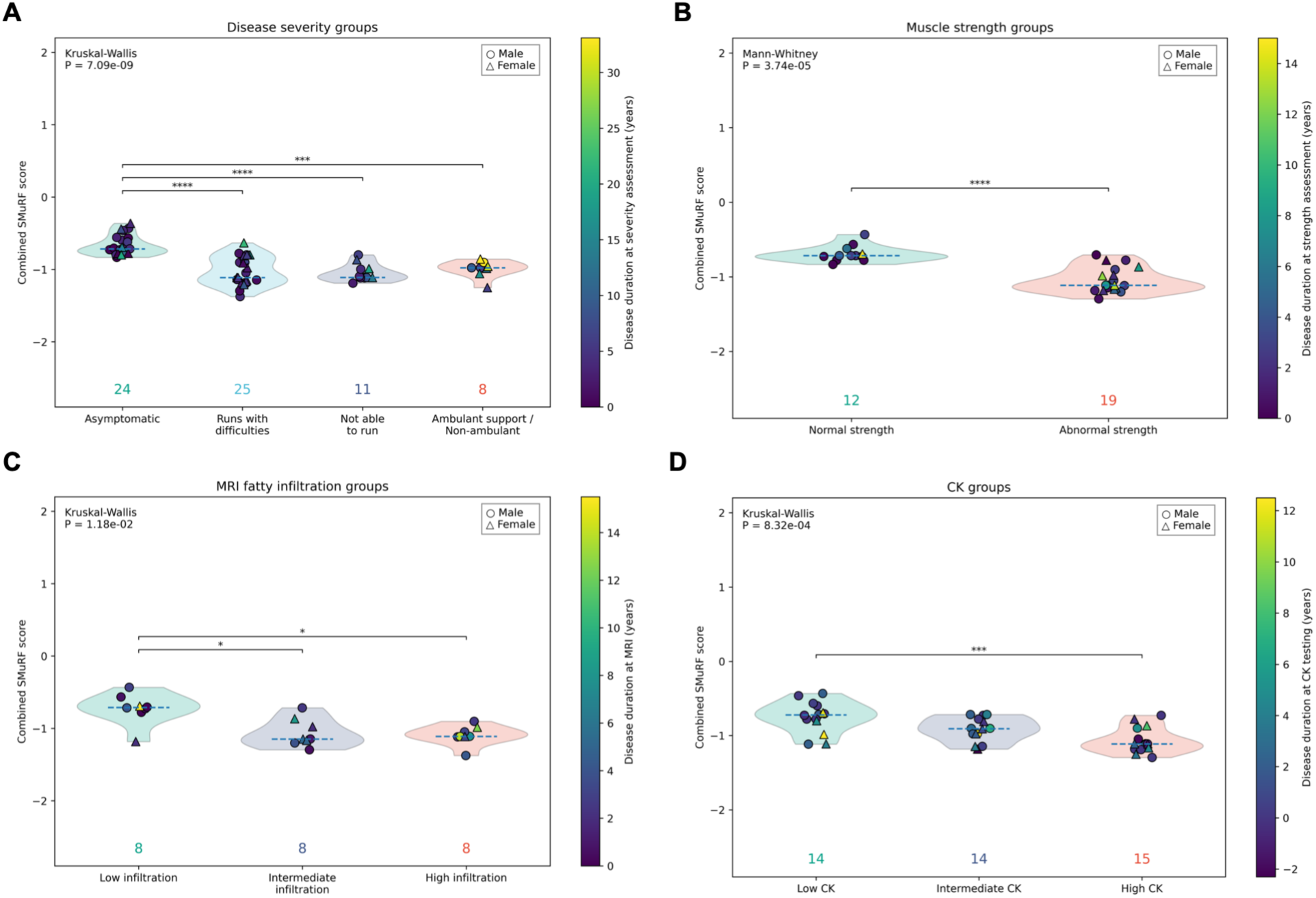
Combined SMuRF scores are associated with clinical disease severity and clinical phenotypes. **(A)** Combined SMuRF scores are associated with disease severity. Combined SMuRF scores were calculated from the SMuRF scores of both alleles. Disease severity was assigned using the most recent available clinical assessment and categorized as: (1) asymptomatic hyperCKemia or minimally affected, (2) ambulatory with difficulty running, (3) ambulatory but unable to run, and (4) ambulatory with support or non-ambulant. **(B)** Combined SMuRF scores are associated with muscle strength. Muscle strength was classified as normal or abnormal based on Medical Research Council (MRC) assessment. **(C)** Combined SMuRF scores are associated with MRI-assessed muscle fatty infiltration. MRI fatty infiltration was categorized into low, intermediate, and high groups based on tertiles of the measured fatty infiltration percentage. **(D)** Combined SMuRF scores are associated with serum creatine kinase (CK) levels. Individuals were stratified into low, intermediate, and high CK groups using tertiles of log10-transformed serum CK levels. Violin plots show the distribution of combined SMuRF scores, and the horizontal dashed blue line indicates the median. Individual symbols represent participants (circles, males; triangles, females), with symbol colors indicating disease duration at the time of the corresponding clinical assessment. Numbers below each violin indicate the number of individuals in each group. Statistical significance was assessed using the Kruskal-Wallis test followed by Dunn’s multiple-comparisons test with Benjamini-Hochberg correction for panels A, C, and D, and a two-sided Mann-Whitney U test for panel B. *P < 0.05; ***P < 0.001; ****P < 0.0001.

To further assess clinical relevance, we examined additional disease-related phenotypes. Patients with abnormal muscle strength exhibited significantly lower combined SMuRF scores than those with normal muscle strength (Figure 5B). Consistent with this observation, higher combined SMuRF scores were associated with greater quantitative muscle strength (Spearman’s ρ = 0.69, p = 1.59e-5, N = 31; Figure S6C). Similarly, patients with intermediate or high muscle fatty infiltration had lower combined SMuRF scores than those with low fatty infiltration (Figure 5C), and combined SMuRF scores were inversely correlated with MRI fatty infiltration (Spearman’s ρ = −0.55, p = 5.67e-3, N = 24; Figure S6D), consistent with reduced disease burden in patients carrying *SGCA* variants with preserved membrane localization. Combined SMuRF scores also decreased across serum creatine kinase (CK) groups (Figure 5D) and were inversely correlated with log10-transformed CK levels (Spearman’s ρ = −0.57, p = 6.89e-5, N = 43; Figure S6E), suggesting that impaired membrane localization is associated with increased muscle membrane instability and ongoing muscle damage. In contrast, combined SMuRF scores were not significantly associated with age at disease onset (Figure S6F). Together, these results demonstrate that experimentally derived SMuRF scores capture clinically meaningful variation in disease severity and correlate with multiple independent measures of disease burden, including muscle strength, muscle fatty infiltration, and serum CK levels, further supporting their utility as functional evidence for *SGCA* variant interpretation.

## Discussion

Accurate interpretation of rare genetic variants remains a major challenge in neuromuscular genetics, particularly for membrane-associated structural proteins such as α-sarcoglycan. Here, we systematically measured the effects of all possible coding SNVs on α-sarcoglycan surface localization using SMuRF^19^, generating a comprehensive functional atlas of *SGCA*. In addition to providing experimental evidence for variant interpretation, our study reveals distinct patterns of functional constraint across α-sarcoglycan, including unexpected properties of the cytoplasmic C-terminal region, highlighting the complexity of variant effects in sarcoglycanopathy.

A major strength of this study is the use of membrane localization as the primary functional readout. Loss of sarcolemmal α-sarcoglycan is a hallmark pathological feature of SGCA-related disease and reflects defects in protein folding, trafficking, complex assembly, or membrane integration.^17,24,25^ By placing an extracellular FLAG epitope immediately downstream of the signal peptide cleavage site, we selectively measured properly processed α-sarcoglycan that successfully reached the cell surface. This design also enabled direct assessment of membrane localization for C-terminal truncating variants that retain the extracellular FLAG epitope. Recent cryo-EM structures of the native rabbit^28^ and mouse^29^ DGCs further support the biological relevance of this assay by revealing that α-sarcoglycan is incorporated into a highly organized extracellular assembly within the SG and DGC complex. All of these support the use of MLSs as an integrated functional readout of α-sarcoglycan maturation and cell-surface localization.

Recent functional studies of *SGCB* demonstrated that experimentally measured sarcolemmal localization is associated not only with variant pathogenicity but also with clinical disease severity. Specifically, patients carrying two severely non-functional *SGCB* alleles experienced significantly earlier loss of ambulation, whereas no association was observed with age at disease onset.^26^ Similarly, combined biallelic SMuRF scores in our SGCA cohort correlated with multiple indicators of disease severity but showed no clear relationship with age at disease onset. The concordance between these independent studies suggests that membrane localization captures a biologically meaningful component of sarcoglycanopathy pathogenesis. Large European cohorts of sarcoglycanopathy patients demonstrated that truncating variants predominantly led to severe phenotypes, while non-truncating variants exhibited a broader phenotypic spectrum which is not easy to predict.^54,55^ In our study, SMuRF scores correlated with clinical severity, suggesting that quantitative functional measurements may help resolve this heterogeneity by distinguishing non-truncating variants with different predicted clinical consequences.

Interestingly, during preparation of this manuscript, the *SGCA* variant A107V, which had previously been classified as benign in ClinVar by the Clinical Genome Resource (ClinGen) Limb Girdle Muscular Dystrophy Variant Curation Expert Panel based primarily on its high population frequency (AF = 0.002741 in East Asian populations in gnomAD v4.1.1), fulfilling the BA1 criterion (AF > 0.002). It was later reclassified as likely pathogenic (April 28, 2026; variation ID: 197094) following the accumulation of clinical evidence from affected individuals.^56–58^ Independently, A107V exhibited a markedly reduced SMuRF Score (−1.46) in our assays (Figure S4B). Despite being enriched in East Asian populations, A107V has been reported in affected individuals of Chinese,^56^ Korean,^57^ and Japanese^58^ ancestry in trans with known pathogenic *SGCA* variants, and affected Japanese individuals carrying A107V have demonstrated reduced expression of sarcoglycan in skeletal muscle, some with limb girdle muscle weakness. Interestingly, A107V was also one of the most frequently observed variants in our Chinese clinical cohort, where 13 of 51 individuals carried the variant, including one homozygous individual (manuscript in preparation). Notably, 9 of these 13 individuals were classified as disease severity category 1, indicating asymptomatic hyperCKemia, or minimal symptoms without functional limitation. Together, these observations suggest that A107V may represent a common Asian founder variant associated with a comparatively mild clinical presentation. Importantly, this example illustrates how discrepancies between experimental functional evidence and existing variant classifications can serve as catalysts for further investigation. In this case, the markedly reduced membrane localization observed in our assay prompted closer examination of the available clinical evidence before the variant was ultimately reclassified by ClinGen. More broadly, systematic functional assays may help prioritize variants whose existing classifications warrant re-evaluation as additional clinical evidence accumulates.

In addition to their clinical utility, experimentally measured SMuRF scores provide an important benchmark for evaluating and potentially improving computational prediction methods. In our previous work on *FKRP* and *LARGE1*, we observed that the computational predictors performed reasonably well overall;^19^ however, this was not the case for *SGCA*. Although predictors such as AlphaMissense and MAVERICK showed moderate concordance with experimental measurements, SMuRF substantially outperformed all computational methods in distinguishing pathogenic from presumed benign variants (Figure 3C). This result is likely because most computational predictors are designed to operate across diverse genes and diseases^59^ and therefore rely primarily on evolutionary conservation, structural features, or machine-learning representations rather than direct measurement of gene-specific biological function.^59,60^ Unlike enzymes such as FKRP, membrane proteins such as SGCA may function in a more complicated manner, concerning membrane localization and complex formation.^47^ Interestingly, MAVERICK was the only computational predictor that partially recapitulated the preservation of SMuRF scores observed for distal C-terminal variants (Figure S4F). Unlike the other predictors evaluated here, MAVERICK was developed for Mendelian disease variant prioritization and incorporates gene-level constraint and population genetic features in addition to protein sequence context.^41^ These additional sources of information may enable it to capture regional patterns of variant tolerance that are not reflected by predictors focused primarily on molecular or evolutionary effects. Nevertheless, MAVERICK still failed to accurately predict many individual variants (Figure 3C and 3E), underscoring the value of experimentally derived functional maps. Our SMuRF dataset may therefore serve as a valuable benchmark for developing improved computational prediction methods for membrane protein-coding genes.

Beyond variant interpretation, the pooled nature of the SMuRF assay also makes it well suited for high-throughput therapeutic screening. Previous studies have demonstrated that several disease-causing *SGCA* missense variants can be rescued by pharmacological approaches, including CFTR correctors, resulting in restoration of sarcolemmal localization and assembly of the SG complex.^31,32^ Future studies could leverage the SMuRF platform to systematically evaluate variant-specific responses to candidate therapeutics, facilitating the identification of rescue-responsive variants and supporting the development of precision medicine strategies for sarcoglycanopathies.

One of the most unexpected findings from this study was the behavior of variants affecting the cytoplasmic C-terminal domain, observed in both the SNV library and nonsense variant library. Unlike variants within the extracellular domains, where both missense and nonsense variants frequently impaired membrane localization, C-terminal missense and nonsense variants consistently retained substantial surface expression, suggesting that the cytoplasmic domain is largely dispensable for α-sarcoglycan trafficking and membrane insertion. This observation is nevertheless difficult to reconcile with human genetic evidence. Several C-terminal nonsense variants, including E342*, E352*, and Q379*, remain exceptionally rare in population databases despite retaining wild-type-like membrane localization (Figure S4D). In addition, two C-terminal nonsense variants, E317* and E352*, have been classified as P or LP in ClinVar, although these classifications appear to rely primarily on predicted loss-of-function mechanisms rather than direct functional or clinical evidence. These observations suggest that although the cytoplasmic tail is largely dispensable for membrane trafficking, disruption of this region may still have biological consequences not captured by membrane localization alone.

A similar separation between membrane localization and biological function has recently been reported for β-dystroglycan,^61^ whose cytoplasmic domain is dispensable for membrane trafficking but remains essential for postsynaptic maturation and neuromuscular function. Recent cryo-EM structures of the rabbit^28^ and mouse^29^ DGCs show that, unlike β-, γ-, and δ-sarcoglycan forming a trimeric scaffold, α-sarcoglycan has a CDHL-SEA architecture, more structurally resembling dystroglycans. The possibility raises that the α-sarcoglycan cytoplasmic tail likewise contributes to functions beyond membrane localization, such as intracellular signaling, mechanotransduction, or regulation of dystrophin-glycoprotein complex stability. Defining these mechanisms will require complementary functional assays that interrogate additional aspects of α-sarcoglycan biology.

An important consideration in interpreting the nonsense library is that the SMuRF assay evaluates the protein-level consequences of truncating variants independently of nonsense-mediated mRNA decay (NMD). In the endogenous genomic context, nonsense variants located within the final exon or within approximately 50-55 nucleotides upstream of the final exon-exon junction are generally predicted to escape NMD.^62–64^ Thus, *SGCA* nonsense variants beginning around residue 319 may produce truncated proteins in vivo rather than undergoing transcript degradation. However, because the SMuRF assay uses *SGCA* CDS constructs, it bypasses the exon junction complex (EJC)-dependent NMD^65^ and therefore directly measures the membrane-localization properties of truncated proteins. In conclusion, the predicted NMD escape and the SMuRF results suggest C-terminal *SGCA* nonsense variants beyond residue 319 may produce truncated proteins that retain membrane localization in vivo.

Several limitations should be considered when interpreting findings in this study. First, as discussed above, the assay specifically measures α-sarcoglycan membrane localization and therefore cannot capture additional pathogenic mechanisms that may occur independently. Moreover, no individuals in our clinical cohort carried C-terminal truncating variants, precluding direct assessment of the clinical consequences of preserved membrane localization. Although two individuals harbored missense variants near our transition zone (R316W and R319K), neither variant provides definitive evidence for this region: R316W yielded a low-confidence SMuRF score (−0.59) and was therefore excluded from downstream analyses, whereas R319K affects the last nucleotide of exon 7 and is predicted to disrupt splicing (SpliceAI Δ = 0.93, donor loss). Because our assay was performed using *SGCA* CDS constructs, pathogenic effects mediated through aberrant RNA splicing are not captured. Furthermore, although key findings were validated in SGCA-deficient human myoblasts, the pooled saturation mutagenesis screen was performed in a heterologous HEK293-based system, which does not fully recapitulate the differentiation state, mechanical environment, or protein interaction landscape of mature skeletal muscle. Finally, although combined SMuRF scores showed strong correlations with multiple clinical measures, validation in larger and more diverse cohorts will further strengthen these findings and increase confidence in their ability to predict clinical severity.

This study provides the first comprehensive functional atlas of *SGCA* coding variation. When integrated with genetic, clinical, and computational evidence, SMuRF scores have the potential to contribute functional evidence for ACMG/AMP variant interpretation, particularly under the PS3 and BS3 criteria. Beyond variant classification, our findings illustrate how comprehensive variant effect maps can reveal biologically meaningful patterns of functional constraint within structural membrane proteins.

## Material and Methods

### Cell Lines and Cell Culture

HEK293T cells (ATCC, CRL-3216; female) were cultured at 37°C in DMEM (Gibco, 11995065) with 10% Fetal Bovine Serum (FBS, R&D Systems, S11150) and 1x Antibiotic-Antimycotic (Anti-anti, Gibco, 15240062). The medium was replaced every 2-3 days. The HEK-BDG stable line was created by co-transduction of Lentiviruses of *SGCB*, *SGCD*, and *SGCG* in HEK293T cells. Monoclonal lines were selected and screened by Western blot to achieve a good co-expression of all 3 proteins. The HEK-BDG cell lines used the same culture conditions as HEK293T cells.

MB135 cells, a healthy human myoblast line,^66^ were cultured at 37°C in Ham’s F-10 Nutrient Mix (Gibco, 11550043) with 20% FBS, 1x Anti-anti, 51 ng/ml dexamethasone (Sigma-Aldrich, D2915), and 10 ng/mL basic fibroblast growth factor (EMD/Millipore, GF003AF-MG). The medium was replaced every 2-3 days. MB135 cells were differentiated in Skeletal Muscle Differentiation Medium (PromoCell, C-23061) with 1x Anti-anti. The differentiation medium was replaced every 4 days. MB135 *SGCA*-KO cell lines used the same culture conditions as MB135 cells.

TrypLE Express Enzyme (Gibco, 12605010) was used to passage cells to maintain cells in healthy confluency (20%-90%). In the scenario of myoblast differentiation into myotubes, cells need to reach 95-100% confluency.

### CRISPR RNP Nucleofection

Synthetic single-guide RNA (sgRNA) was designed in-house and ordered from Integrated DNA Technologies, and SpCas9 2NLS nuclease was purchased from Synthego. RNP complexes were prepared in SE Cell Line Nucleofector Solution (Lonza, PBC1-00675) and delivered into cells with a Lonza 4D-Nucleofector. The program used for MB135 was CA-137. Monoclonal lines were isolated from pooled nucleofected cells and genotyped by targeted Sanger sequencing. sgRNA sequences and genotyping primers are provided in Table S2. MB135 *SGCA*-KO monoclonal line A2 carries a homozygous 1-bp insertion (NM_000023.4: c.92dup, insert T).

### Plasmid Construction

Lenti-*SG gene* plasmids used the backbone of Lenti-UbC-FKRP-EF1a-BSD (Addgene, #205150).^19,67^ Coding sequences (CDS) of *SGCA* (NM_000023.4), *SGCB* (NM_000232.5), *SGCD* (NM_000337.6), and *SGCG* (NM_000231.3) were cloned from cDNA generated from RNA isolated from MB135 myoblast-differentiated myotubes. The EF-1α core promoter was used to drive *SG gene* expression. A linker and FLAG tag were inserted right after the single peptide of SGCA (c.1-69), followed by the rest of *SGCA* cDNA (c.70-1161). A linker plus a 6xHis, c-Myc, or V5 tag was added after *SGCB*, *SGCD*, or *SGCG* cDNA, respectively, followed by a stop codon. For *SGCB*, *SGCD*, and *SGCG* constructs, the drug resistance marker was a BSD driven by IRES2; for the *SGCA* construct, a PuroR driven by an hPGK promoter was used. BSD encodes blasticidin S deaminase, which confers blasticidin resistance. PuroR encodes puromycin N-acetyltransferase, which confers puromycin resistance. Plasmid assemblies were achieved either with NEBuilder HiFi DNA Assembly Master Mix (NEB, E2621) or T4 DNA Ligase (M0202). Plasmid sequences are listed in Table S2. The introduction of individual variants to the lentiviral plasmids used Site-Directed Mutagenesis by PCR.

### Lentiviral Packaging and Transduction

Lentivirus was packaged in HEK293T cells. For a 10-cm dish of HEK293T cells (90% confluency), 10 μg psPAX2 (Addgene, #12260), 2 μg pMD2.G (Addgene, #12259), and 9 μg lentiviral plasmid pool were mixed in 1.5 mL Opti-MEM (Gibco, 31985062), then 45 μL TransIT-LT1 Transfection Reagent (Mirus, MIR 2300) was added. The preparation was gently mixed, incubated at room temperature for 15 min, and then added to the cells dropwise. The volume was adjusted to 17 mL with culture medium, and the dish was gently moved in a cross-shaped pattern to ensure even distribution. 72 hrs later, the supernatant in the dish was filtered with a 0.45 μm PES filter (Thermo Scientific, 165-0045), mixed with 5 mL Lenti-X Concentrator (Takara, 631232), and rocked at 4 °C overnight. The viral particles were then spun down (1,800 × g, 4 °C, 1hr) and resuspended in 200 μL DMEM. Lentivirus was titrated with Lenti-X GoStix Plus (Takara, 631280).

For lentiviral transduction, the cells to be transduced were plated in wells of plates. One day after seeding, remove 200 μL of medium from each well, and replace with fresh culture media supplemented with polybrene (Sigma-Aldrich, 107689; final concentration 8 μg/mL) and lentiviruses, for a spinfection (800 × g, 32 °C, 1hr). One day post-transduction, the medium was replaced, and drug selection was started if applicable. For constructs with BSD, Blasticidin S HCl (Gibco, A1113903, final conc. 10 μg/mL) was used for drug selection. For constructs with PuroR, Puromycin Dihydrochloride (Gibco, A1113803, final conc. 1-10 μg/mL) was used. Drug selection was performed for 7-14 days.

### *SGCA* Saturation Mutagenesis Library Design and Construction

For the SNV library, each possible SNV across the *SGCA* CDS was individually encoded in a 65-nt single-stranded DNA (ssDNA) oligonucleotide. For the nonsense variant library, each codon spanning the C-terminal 150 amino acids (residues 238-387) of the *SGCA* CDS was individually replaced with a TAA stop codon and encoded in a 67-nt ssDNA oligonucleotide. The oligonucleotides were synthesized as a pooled library by Twist Bioscience and cloned using programmed allelic series with common procedures cloning (PALS-C), following the SMuRF workflow described in our previous work^19,67^. Briefly, oligos were used to introduce programmed variants into block-specific *SGCA* fragments by PCRs, which were subsequently assembled into the lentiviral expression backbone using NEBuilder HiFi DNA Assembly (NEB, E2621). The library design ensured compatibility with short-read NGS, enabling inclusion of synonymous, missense, nonsense, and start-loss variants while minimizing synthesis cost and amplification bias. The final plasmid pools were transformed into Endura electrocompetent cells (Lucigen, 60242-1) via electrotransformation (Bio-Rad Gene Pulser II) and expanded for plasmid preparation. Library complexity was assessed by colony-forming unit counts, with at least 110× theoretical coverage achieved for each *SGCA* block.

### Plasmid Library QC and Lentivirus Library Production

Quality control (QC) of the plasmid sub-pools was performed using the Amplicon-EZ service from GENEWIZ to assess variant representation and distribution within each sub-pool (Figures S1C and S1D). Lentivirus of each block was packaged by HEK293T cells in one 10-cm dish per block within the same experiment batch. Lentiviral input was scaled for each block based on the GoStix value (GV) measured using Lenti-X GoStix Plus (Takara, 631280), with 1e4 GV·μL of lentivirus used per block. For transduction, 720k HEK-BDG cells were plated in one well of a 6-well plate for each block, with cell numbers determined using an automated cell counter (Bio-Rad, TC20). Cells from each block were subsequently expanded to more than 30 million cells for FACS.

### Live-Cell Surface Staining and FACS

Block 2 of the *SGCA* library was used to evaluate and optimize FACS separation for the full library. The optimized staining and FACS workflow for one representative block is described here. After medium removal, cells were washed once with DPBS (Gibco, 14190144), detached with 6 mL Versene (Gibco, 15040066) per T175 flask at 37°C for 15 min, and counted. For each block, 30 million cells were transferred to a 15-mL tube for staining. Cells were pelleted at 600 × g for 10 min at 4°C, washed once with 3 mL DPBS, and pelleted again at 600 × g for 5 min at 4°C. Cells were then resuspended in 1 mL cold sorting buffer containing 0.5% bovine serum albumin (BSA) and 1 mM EDTA in DPBS, supplemented with 2.5 μL anti-FLAG antibody APC conjugate (BioLegend, 637308) and 3.3 μL Viobility 405/452 Fixable Dye (Miltenyi Biotec, 130-130-420). Sorting buffer was prepared by mixing 1 volume MACS BSA Stock Solution (Miltenyi Biotec, 130-091-376), 0.04 volume 0.5 M EDTA (AmericanBio, AB00502-01000), and 19 volumes DPBS. All subsequent steps were performed protected from light. Samples were gently rocked on ice for 30 min in a cold room, after which 7 mL sorting buffer was added. Cells were pelleted at 600 × g for 10 min at 4°C, resuspended in 3 mL sorting buffer, and filtered through a 40-μm cell strainer (Falcon, 352340).

Fluorescence flow cytometry experiments were performed on a BD LSR II flow cytometer to evaluate staining signal before sorting. FACS experiments were performed on a BD FACSAria II cell sorter. Forward scatter (FSC) and side scatter (SSC) were used to exclude debris and multiplets, and singlets were retained for downstream analysis. BV421 (450/50 bandpass filter) was used to detect Viobility 405/452 Fixable Dye and gate live cells. APC (670/30 bandpass filter) was used to detect APC signal from the anti-FLAG antibody conjugate. For each block, at least 20,000 events were recorded to determine gating parameters. For FACS, the top ~20% of cells were collected as the high-surface-expression fraction, and the bottom ~30% were collected as the low-surface-expression fraction. A minimum coverage of ~1,500× was achieved for each sorted fraction of each block, corresponding, for example, to more than 1.1 million cells collected per fraction for a block containing 717 variants.

### Genomic DNA Extraction, Library Preparation, and Sequencing

Sorted cells were pelleted at 800 × g for 10 min at 4°C, and genomic DNA was extracted from each sample using the PureLink Genomic DNA Mini Kit (Invitrogen, K182002). Sequencing libraries were prepared using a three-step PCR workflow. In the first PCR, lentiviral CDS sequences were amplified from each sample using primers specific to the lentiviral backbone. In the second PCR, block-specific regions were isolated using primers containing a 20-bp block-specific flanking sequence and a partial Illumina adapter sequence. Forward primers also included barcodes to distinguish the high- and low-surface-expression fractions. In the third PCR, the remaining Illumina adapter sequences were added to the amplicons. Final PCR products were pooled equimolarly and submitted to short-read paired-end NGS. SNV libraries were sequenced by Psomagen (Illumina NovaSeq X Plus 10B, 2 × 150 bp), yielding approximately 1.25 billion paired-end reads per library. Nonsense libraries were sequenced by Ampseq (Avitia platform, 2 × 300 bp), yielding approximately 650,000 paired-end reads per library. The three-step PCR primers are provided in Table S2.

### Variant Counting and *SGCA* SMuRF Score Generation

Raw sequencing data were processed using the “Gargamel” pipeline.^19^ Briefly, reads containing more than one variant were removed, and variant counts were calculated for each sorted fraction. For each variant, enrichment was calculated in the high- and low-surface-expression fractions, and relative enrichment was normalized to the corresponding wild-type sequence within each block to generate replicate-level MLSs. Variant counts from three biological replicates were then integrated using DiMSum^33^ to generate combined variant-level scores and associated error estimates. Scores were normalized by block using synonymous variants as neutral references and converted to log2 scale to generate the final *SGCA* SMuRF scores. Variant-level uncertainty was estimated by DiMSum as sigma (σ). For confidence filtering, the σ distribution of synonymous variants, rather than that of all variants, was used as the reference distribution because synonymous variants are expected to be functionally neutral and therefore more accurately reflect replicate-level technical and biological variation. Because the *SGCA* dataset had higher overall coverage and exhibited a narrower σ dynamic range than the previously reported SMuRF-*FKRP* and *LARGE1* datasets, the previously used Q3 + 1.5 × interquartile range (IQR) threshold was considered overly stringent for this dataset. Therefore, a modified threshold of Q3 + 3.0 × IQR based on the synonymous-variant σ distribution was used to identify low-confidence variants. Variants exceeding this threshold were classified as low confidence and excluded from downstream analyses.

### Nonsense Variant Counting and Score Generation

Raw sequencing data from the nonsense library were processed using the “nonsense_ library” pipeline under “DeniSa”. For each biological replicate, reads containing a single TAA codon were identified and assigned to the corresponding *SGCA* codon. Variant counts were determined separately for the high- and low-surface-expression fractions. To account for differences in sequencing depth, a pseudocount of 1 was added to each variant count, and normalized variant frequencies were calculated for each fraction. Variant enrichment was then calculated as the log2 ratio of normalized frequencies between the high- and low-surface-expression fractions. Final mean nonsense scores (Figure 2E) were calculated as the mean log2 enrichment score from two independent biological replicates. Replicate reproducibility was assessed by Pearson correlation. Nonsense variants were classified as low confidence if the combined read count (high-and low-surface-expression fractions) was fewer than 100 in either replicate or if the inter-replicate standard deviation exceeded 1.5.

### Clinical Annotation and Population Frequency Analysis

Clinical variant annotations were retrieved from ClinVar. The version we downloaded was from February 2^nd^, 2026 (Figure S4B only) and June 14^th^, 2026 (all other figures). Variants classified as P, P/LP, LP, B, or LB were used to benchmark SMuRF scores.

Population allele frequencies were obtained from the exome sequencing database at the Genome Aggregation Database (gnomAD v4 exomes). Variants were grouped by allele frequency to assess enrichment of deleterious functional scores among rare variants and convergence toward wild-type scores among common variants.

### Comparison of SMuRF Score to *in silico* Predictors

To compare the performance of *SGCA* SMuRF scores with computational predictors, receiver operating characteristic (ROC) curve analysis was performed following the framework described in the SMuRF study,^19^ with additional predictors included. The analysis was restricted to missense variants scored by SMuRF and all evaluated predictors to enable a matched comparison. The positive reference set comprised ClinVar missense variants classified as P, P/LP, or LP. The negative reference set consisted of common missense variants observed in gnomAD v4 exomes (AF > 1e-5) and not classified as P, P/LP, or LP in ClinVar. Computational predictors included AlphaMissense, MAVERICK, REVEL, MetaSVM, VESM3B, MutScore, PrimateAI-3D, CADD, EVE, and MVP. ROC curves were calculated using the pROC package in R. Sensitivity and specificity were calculated across score thresholds for each method, and AUC was used as the summary measure of discriminatory performance.

### Calculation of Combined SMuRF Scores for Biallelic Genotypes

To estimate the combined effect of biallelic *SGCA* variants, individual SMuRF scores were integrated into a combined SMuRF score representing the predicted overall membrane localization capacity contributed by both alleles. Because SMuRF scores are reported on a log₂ scale, scores were first transformed into linear space, summed across alleles, and then converted back to log₂ space according to the following equation:

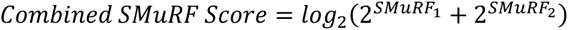

where SMuRF₁ and SMuRF₂ represent the experimentally determined SMuRF scores of the two *SGCA* alleles. This approach assumes additive contributions of the two alleles to total α-sarcoglycan membrane localization capacity.

Patients carrying two non-SNV pathogenic variants were excluded from analyses requiring combined SMuRF scores because neither allele could be directly assigned a SMuRF score. Patients carrying at least one *SGCA* CDS SNV with a SMuRF score were retained. For pathogenic variants lacking SMuRF scores, including small insertions/deletions, copy-number variants, and splice-altering variants, a surrogate SMuRF score of −2.18 was assigned. This value was derived from the median SMuRF score of ClinVar P, P/LP, and LP variants after excluding the two outlier pathogenic variants R110L and E352* in this assay (Figure S4B), as well as splice-region variants affecting the first or last nucleotide of exons that are expected to exert pathogenicity primarily through aberrant splicing rather than altered protein localization (column “splicing-region variant” in Table S1). For the pathogenic variant c.233_234delinsGA (p.Tyr78Ter), identified in seven patients (one homozygous and six compound heterozygous) in our Chinese cohort, although this variant was not directly represented in our SNV library, it produces the same protein consequence as the SNVs c.234C>A (p.Tyr78Ter) (SMuRF score = −2.15) and c.234C>G (p.Tyr78Ter) (SMuRF score = −2.35). Therefore, its SMuRF score was assigned as the mean of SMuRF scores for these two variants (−2.25). Variants with low-confidence SMuRF scores were excluded from combined score analyses. Clinical severity analyses were subsequently performed using the resulting combined SMuRF scores.

### Clinical Cohort and Phenotype Analyses

A clinically characterized cohort of 68 deidentified individuals with *SGCA*-associated disease was assembled through collaborating neuromuscular centers in China and the United States. Clinical data from the Chinese cohort (51 patients in total) were provided by collaborating neuromuscular centers, whereas U.S. participants (17 patients) were identified through the Global Rare Disease Patient Registry Data Repository (GRASP-LGMD) following an approved data request and were required to carry two pathogenic or likely pathogenic *SGCA* variants. For each individual, a combined SMuRF score was calculated using the formula described above (Table S3).

Clinical disease severity was categorized into four groups: (1) asymptomatic hyperCKemia or minimally affected, (2) ambulatory with difficulty running, (3) ambulatory but unable to run, and (4) ambulatory with support or non-ambulant. For participants from the Chinese cohort, clinical disease severity was assigned by clinicians using the most recent available clinical assessment. For GRASP participants, disease severity was assigned through review of ambulatory status and the most recent North Star Assessment for Limb-Girdle Type Muscular Dystrophies (NSAD), with particular emphasis on the Walk10m and Run10m items. Ambulatory status was prioritized when discrepancies were present among clinical measures.

Analyses of muscle strength, MRI fatty infiltration, and serum creatine kinase (CK) were performed using the subset of participants from the Chinese cohort for whom these data were available. Muscle strength was evaluated using the Medical Research Council (MRC) scale across 13 muscle groups, including neck flexion, shoulder abduction/adduction, elbow flexion/extension, hand grip, hip flexion/abduction/adduction, knee flexion/extension, and ankle dorsiflexion/plantar flexion. Individuals were classified as having normal muscle strength if all evaluated muscle groups had an MRC grade of 5; the presence of an MRC grade < 5 in any muscle group was classified as abnormal muscle strength. For correlation analyses, a quantitative muscle strength score was calculated as the percentage of the maximum possible total MRC score across all evaluated muscle groups. MRI fatty infiltration was quantified as the summed fatty infiltration score across 13 thigh muscles (0-5 points per muscle), normalized to the maximum possible score (13 × 5) and expressed as a percentage. Participants were subsequently stratified into low, intermediate, and high MRI fatty infiltration groups using tertiles of the percentage values. Serum CK levels (IU/L) were log10-transformed for continuous analyses, and participants were stratified into low, intermediate, and high CK groups using tertiles of the log10-transformed values. Disease duration was calculated as the age at phenotypic assessment minus the age at symptom onset. Negative disease duration values were permitted for serum CK analyses because some CK measurements were obtained before symptom onset.

For categorical comparisons, combined SMuRF scores were compared across disease severity, MRI fatty infiltration, and CK groups using the Kruskal-Wallis test followed by Dunn’s multiple-comparisons test with Benjamini-Hochberg correction. Muscle strength groups were compared using two-sided Mann-Whitney U tests. Associations between combined SMuRF scores and continuous clinical variables, including disease severity, muscle strength, MRI fatty infiltration, serum CK levels, and age at symptom onset, were assessed using Spearman’s rank correlation coefficient. Within each disease severity group, male and female participants were compared using two-sided Mann-Whitney U tests, with P values adjusted using the Benjamini-Hochberg procedure.

### Western Blotting

Cells were lysed in ice-cold radioimmunoprecipitation assay (RIPA) lysis and extraction buffer (Pierce, PI89900) supplemented with protease inhibitor cocktail (Sigma-Aldrich, P8340-1ML; 10 μL per 1 mL RIPA buffer). Samples were rocked on ice for 15 min and centrifuged at 16,000 × g for 30 min at 4°C. The supernatant was transferred to a clean 1.5-mL microcentrifuge tube for downstream analysis.

Protein concentration was measured using the Bio-Rad DC Protein Assay II kit (Bio-Rad, 5000112) according to the manufacturer’s instructions. BSA (Sigma-Aldrich, A9647) standards ranging from 0 to 1.5 mg/mL were prepared in the same lysis buffer, and absorbance at 750 nm was measured using a plate reader (Bio-Rad, 1681135). Protein concentrations were calculated from the BSA standard curve.

For each sample, approximately 10 μg of protein was adjusted to an equal volume using lysis buffer. Four-fold Laemmli sample buffer (Bio-Rad, 1610747) supplemented with 10% β-mercaptoethanol was added at a 1:3 ratio to each sample. Samples were briefly vortexed, heated at approximately 95°C for 5 min, cooled on ice, and briefly centrifuged. Samples and 3.5 μL PageRuler Plus Prestained Protein Ladder (Thermo Fisher Scientific, 26619) were loaded onto 4-15% Mini-PROTEAN TGX stain-free protein gels (Bio-Rad, 4568084). SDS-PAGE was performed using a Mini-PROTEAN Tetra Cell system (Bio-Rad, 1658000) at 120 V for approximately 60 min. Proteins were transferred to nitrocellulose membranes (Bio-Rad, 1704158) using the Trans-Blot Turbo system (Bio-Rad, 1704150EDU) with the 7-min transfer program.

Vinculin was used as a loading control, and membranes were cut at the appropriate position to separately probe sarcoglycan proteins and vinculin. MyHC was used to assess myogenic differentiation in myoblast-derived myotube samples. Membranes were blocked for 1 h at room temperature in blocking buffer consisting of 5% OmniBlock non-fat dry milk (AmericanBio, AB10109-00100) in 1× TBS-Tween 20 (TBS-T). Primary antibodies were diluted in blocking buffer and incubated with membranes overnight at 4°C with rocking. The following primary antibodies were used: anti-vinculin (Sigma-Aldrich, V9131, 1:80,000), anti-MyHC (Invitrogen, 14-6503-37, 1:500), anti-SGCA (Invitrogen, MA5-37982, 1:1,000), anti-SGCB (MilliporeSigma, HPA011422, 1:200), anti-SGCD (abcam; ab137101, 1:1,000), and anti-SGCG (GeneTex, GTX117176, 1:2,000).

The next day, membranes were washed five times for 5 min each in TBS-T and incubated with HRP-conjugated secondary antibodies for 2 h at room temperature. Anti-rabbit HRP-conjugated secondary antibody (Cell Signaling Technology, 7074S, 1:2,000) and anti-mouse HRP-conjugated secondary antibody (Cell Signaling Technology, 7076S, 1:2,000) were diluted in blocking buffer. Membranes were then washed five times for 5 min each in TBS-T and developed using Clarity Max ECL substrate (Bio-Rad, 1705062) or Clarity ECL substrate (Bio-Rad, 1705060). Chemiluminescent and colorimetric images were acquired using a ChemiDoc imaging system (Bio-Rad, 12005606).

### Immunofluorescence

For myoblast-differentiated myotube staining, 15-mm round Thermanox coverslips (Thermo Scientific, 174969) were placed in 24-well plates and coated with 0.1% gelatin (Sigma-Aldrich, G9391) by a transient rinse. After air-drying, MB135-derived cells were seeded onto coverslips in 0.7 mL growth medium at approximately 40% confluency. Five days after plating, when cells reached 100% confluency, growth medium was replaced with differentiation medium, and cells were differentiated for 3-4 days until mature myotubes formed. Cells were washed once with DPBS and fixed with 4% paraformaldehyde (PFA; Sigma-Aldrich, 158127) in DPBS for 10 min at room temperature. Cells were blocked with 2% BSA (Sigma-Aldrich, A9647) in DPBS for 1 h at room temperature. Primary antibodies were diluted in 1% BSA and incubated overnight at 4°C in a humidified chamber. The following primary antibodies were used: anti-MyHC (Invitrogen, 14-6503-37, 1:500), anti-SGCA (Invitrogen, MA5-37982, 1:400), and anti-SGCA (Leica, NCL-L-a-SARC-A, 1:30). Cells were washed three times with DPBS and incubated with secondary antibodies diluted in 1% BSA for 2 h at room temperature in the dark. The following secondary antibodies were used: goat anti-mouse IgG Alexa Fluor 488 (Invitrogen, A32723, 1:300) and goat anti-rabbit Alexa Fluor 555 (Invitrogen, A32732, 1:300). Coverslips were washed three times with DPBS and mounted cell-side down onto microscope slides (Fisher Scientific, 22-037-246) using VECTASHIELD HardSet Antifade Mounting Medium with DAPI (Vector Laboratories, H-1500-10). Slides were kept at room temperature for 20 min in the dark before imaging.

For HEK293T-derived cell staining, cells were seeded in Nunc Lab-Tek chamber slides (Thermo Fisher Scientific, 177399) coated with 50 μg/mL poly-D-lysine (Gibco, A3890401) in DPBS. For coating, 400 μL poly-D-lysine was added to each well, incubated for 1 h at room temperature in a sterile hood, washed three times with sterile Milli-Q water, and air-dried. HEK293T-derived cells were seeded in 0.7 mL growth medium at approximately 50% confluency. The following day, chamber slides were placed on ice at 4°C for 30 min, medium removed afterwards, and gently washed with ice-cold DPBS once. For live-cell surface staining, cells were incubated with primary antibodies for 2 h on ice at 4°C, gently washed three times with ice-cold DPBS, and incubated with secondary antibodies for 2 h on ice at 4°C in the dark. Primary antibodies included anti-FLAG (Cell Signaling Technology, 14793, 1:400), anti-6x-His tag (Invitrogen, MA1-21315, 1:1,000), anti-c-Myc tag (Invitrogen, MA1-980, 1:250), and anti-V5 tag (Abcam, ab9116, 1:300). Secondary antibodies included goat anti-rabbit IgG Alexa Fluor 488 (Invitrogen, A-11008, 1:300) and goat anti-mouse IgG Alexa Fluor 488 (Invitrogen, A32723, 1:300). Both primary and secondary antibodies were diluted in DPBS containing 0.5% BSA. Cells were then washed twice with DPBS, fixed with 4% PFA for 10 min at room temperature, washed twice with DPBS, and mounted with VECTASHIELD HardSet Antifade Mounting Medium with DAPI before applying coverslips (Globe Scientific, 1414-10) and sealing with clear nail polish. Slides were kept at room temperature for 20 min in the dark before imaging.

Images were acquired using a Revolve ECHO microscope with DAPI, FITC, and Texas Red channels.

### Structural Visualization and Residue-Level Score Mapping

Protein 3D structures from the Protein Data Bank (PDB) were visualized using UCSF ChimeraX v1.11.1. The cryo-EM structures of the native DGC from rabbit (PDB: 9C3C; Liu et al., 2025)^28^ and mouse (PDB: 8YT8; Wan et al., 2025)^29^ were used. The AlphaFold-predicted human SGCA structure was obtained from the AlphaFold Protein Structure Database (UniProt Q16586). Figures displaying domain organization or residue-level mean missense scores on 3D structures were generated using custom ChimeraX command files. The residue-level mean missense score was calculated as the mean SMuRF score of all high-confidence missense substitutions at each amino acid position. Structural domains were annotated according to the SGCA linear protein sequence and adapted from the rabbit structure described by Liu et al.^28^

Human SGCA residue-level scores were mapped onto the SGCA of the native rabbit DGC structure. Only residues resolved in the structure were displayed. Based on protein sequence alignment (Figure S5C), rabbit SGCA residue 1 corresponds to human residue 1 but was not resolved in PDB 9C3C; rabbit residue 2 is an additional residue relative to human SGCA and was also not resolved; rabbit residues 3-54 correspond to human residues 2-53; therefore, human residue-level scores for residues 2-53 were mapped to rabbit residues 3-54 in the 3D structure. Human residue 54 is absent from the rabbit sequence and was therefore not displayed. Rabbit residues 55-387 correspond to human residues 55-387, and human residue-level scores for this region were mapped directly to the corresponding rabbit residues. For mouse SGCA and the AlphaFold-predicted human SGCA structure, human residue-level scores were directly mapped to the corresponding residues (Figures S5A and S5B).

### Statistical Analysis

For functional assay analyses, two-sided Wilcoxon tests were performed with the “ggsignif” R package. Spearman’s rank correlation coefficients were calculated with the “cor.test” function in R.

For clinical analyses, statistical analyses were performed using Python 3. Spearman’s rank correlation coefficients were calculated using the “spearmanr” function from SciPy. Comparisons between two clinical groups were performed using two-sided Mann-Whitney U tests, whereas comparisons among three or more groups were performed using Kruskal-Wallis tests followed by Dunn’s multiple-comparison test with Benjamini-Hochberg false discovery rate correction. Sex-stratified comparisons were adjusted for multiple testing using the Benjamini-Hochberg procedure.

## Acknowledgments

S.H. is supported by a Muscular Dystrophy Association Development Grant (MDA 963708) for “Improving interpretation of variants in Sarcoglycanopathies”. This work is supported in part by funding from Cure LGMD2D Research Foundation and LGMD2D Foundation. M.L. is supported by the Chan Zuckerberg Initiative Science Diversity Leadership Award (2022-309740). We thank Nicholas E. Johnson and the GRASP LGMD consortium for granting us access to their deidentified patient data. We thank Stephen Tapscott for the gift of MB135 cells.

## Author Contributions

S.H., K.M., and M.L. conceived and designed this study. M.L. supervised the experiments and analyses of this study. S.H. designed and performed the experiments for the establishment, application, and validation of SMuRF *SGCA*, with K.M.’s help. A.L. and Y.Y. assisted with plasmid library electrotransformation. K.K.N., K.M., S.H., and M.L. developed the analytical pipeline. K.K.N., S.H., and M.L. performed computational analyses with K.M.’s help. Y.L., Z.X., C.H., B.Z., and W.Z. provided deidentified clinical data. S.H., K.K.N., K.M., S.K., and M.L. wrote this manuscript with input from other authors.

## Data and Code Availability

The sequencing data generated in this study have been deposited in the NCBI Sequence Read Archive under BioProject accession PRJNA1472910. The associated BioSample accessions are SAMN60524398, SAMN60524399, and SAMN60524400. The associated SRA run accessions are SRR38916725, SRR38916726, and SRR38916727. Data-processing pipelines and analysis code are publicly available at https://github.com/leklab/Gargamel and https://github.com/leklab/DeniSa. The variant scores have been uploaded to MaveDB (urn:mavedb:00001283-a-1).

## References

1. Duggan, D.J., Gorospe, J.R., Fanin, M., Hoffman, E.P., and Angelini, C. (1997). Mutations in the sarcoglycan genes in patients with myopathy. N. Engl. J. Med. 336, 618–624.

2. Politano, L., Nigro, V., Passamano, L., Petretta, V., Comi, L.I., Papparella, S., Nigro, G., Rambaldi, P.F., Raia, P., Pini, A., et al. (2001). Evaluation of cardiac and respiratory involvement in sarcoglycanopathies. Neuromuscul. Disord. 11, 178–185.

3. Ozawa, E., Mizuno, Y., Hagiwara, Y., Sasaoka, T., and Yoshida, M. (2005). Molecular and cell biology of the sarcoglycan complex. Muscle Nerve 32, 563–576.

4. Gumerson, J.D., and Michele, D.E. (2011). The dystrophin-glycoprotein complex in the prevention of muscle damage. J. Biomed. Biotechnol. 2011, 210797.

5. Gao, Q.Q., and McNally, E.M. (2015). The dystrophin complex: structure, function, and implications for therapy. Compr. Physiol. 5, 1223–1239.

6. Wilson, D.G.S., Tinker, A., and Iskratsch, T. (2022). The role of the dystrophin glycoprotein complex in muscle cell mechanotransduction. Commun. Biol. 5, 1022.

7. Moreira, E.S., Vainzof, M., Suzuki, O.T., Pavanello, R.C.M., Zatz, M., and Passos-Bueno, M.R. (2003). Genotype-phenotype correlations in 35 Brazilian families with sarcoglycanopathies including the description of three novel mutations. J. Med. Genet. 40, E12.

8. Fanin, M., Duggan, D.J., Mostacciuolo, M.L., Martinello, F., Freda, M.P., Sorarù, G., Trevisan, C.P., Hoffman, E.P., and Angelini, C. (1997). Genetic epidemiology of muscular dystrophies resulting from sarcoglycan gene mutations. J. Med. Genet. 34, 973–977.

9. Duggan, D.J., Manchester, D., Stears, K.P., Mathews, D.J., Hart, C., and Hoffman, E.P. (1997). Mutations in the delta-sarcoglycan gene are a rare cause of autosomal recessive limb-girdle muscular dystrophy (LGMD2). Neurogenetics 1, 49–58.

10. Moore, S.A., Shilling, C.J., Westra, S., Wall, C., Wicklund, M.P., Stolle, C., Brown, C.A., Michele, D.E., Piccolo, F., Winder, T.L., et al. (2006). Limb-girdle muscular dystrophy in the United States. J. Neuropathol. Exp. Neurol. 65, 995–1003.

11. 11. Carrié, A., Piccolo, F., Leturcq, F., de Toma, C., Azibi, K., Beldjord, C., Vallat, J.M., Merlini, L., Voit, T., Sewry, C., et al. (1997). Mutational diversity and hot spots in the alpha-sarcoglycan gene in autosomal recessive muscular dystrophy (LGMD2D). J. Med. Genet. 34, 470–475.

12. Landrum, M.J., Lee, J.M., Riley, G.R., Jang, W., Rubinstein, W.S., Church, D.M., and Maglott, D.R. (2014). ClinVar: public archive of relationships among sequence variation and human phenotype. Nucleic Acids Res. 42, D980–5.

13. Mendell, J.R., Rodino-Klapac, L.R., Rosales-Quintero, X., Kota, J., Coley, B.D., Galloway, G., Craenen, J.M., Lewis, S., Malik, V., Shilling, C., et al. (2009). Limb-girdle muscular dystrophy type 2D gene therapy restores alpha-sarcoglycan and associated proteins. Ann. Neurol. 66, 290–297.

14. Mendell, J.R., Chicoine, L.G., Al-Zaidy, S.A., Sahenk, Z., Lehman, K., Lowes, L., Miller, N., Alfano, L., Galliers, B., Lewis, S., et al. (2019). Gene Delivery for Limb-Girdle Muscular Dystrophy Type 2D by Isolated Limb Infusion. Hum. Gene Ther. 30, 794–801.

15. Pejaver, V., Byrne, A.B., Feng, B.-J., Pagel, K.A., Mooney, S.D., Karchin, R., O’Donnell-Luria, A., Harrison, S.M., Tavtigian, S.V., Greenblatt, M.S., et al. (2022). Calibration of computational tools for missense variant pathogenicity classification and ClinGen recommendations for PP3/BP4 criteria. Am. J. Hum. Genet. 109, 2163–2177.

16. Bergquist, T., Stenton, S.L., Nadeau, E.A.W., Byrne, A.B., Greenblatt, M.S., Harrison, S.M., Tavtigian, S.V., O’Donnell-Luria, A., Biesecker, L.G., Radivojac, P., et al. (2025). Calibration of additional computational tools expands ClinGen recommendation options for variant classification with PP3/BP4 criteria. Genet. Med. 27, 101402.

17. Sandonà, D., and Betto, R. (2009). Sarcoglycanopathies: molecular pathogenesis and therapeutic prospects. Expert Rev. Mol. Med. 11, e28.

18. Ma, K., Gauthier, L.O., Cheung, F., Huang, S., and Lek, M. (2024). High-throughput assays to assess variant effects on disease. Dis. Model. Mech. 17,.

19. Ma, K., Huang, S., Ng, K.K., Lake, N.J., Joseph, S., Xu, J., Lek, A., Ge, L., Woodman, K.G., Koczwara, K.E., et al. (2024). Saturation mutagenesis-reinforced functional assays for disease-related genes. Cell 187, 6707–6724.e22.

20. Ervasti, J.M., and Campbell, K.P. (1991). Membrane organization of the dystrophin-glycoprotein complex. Cell 66, 1121–1131.

21. Straub, V., and Campbell, K.P. (1997). Muscular dystrophies and the dystrophin-glycoprotein complex. Curr. Opin. Neurol. 10, 168–175.

22. Kirschner, J., and Lochmüller, H. (2011). Sarcoglycanopathies. Handb. Clin. Neurol. 101, 41–46.

23. Zanotti, S., Magri, F., Poggetti, F., Ripolone, M., Velardo, D., Fortunato, F., Ciscato, P., Moggio, M., Corti, S., Comi, G.P., et al. (2022). Immunofluorescence signal intensity measurements as a semi-quantitative tool to assess sarcoglycan complex expression in muscle biopsy. Eur. J. Histochem. 66,.

24. Holt, K.H., and Campbell, K.P. (1998). Assembly of the sarcoglycan complex. Insights for muscular dystrophy. J. Biol. Chem. 273, 34667–34670.

25. Draviam, R.A., Wang, B., Shand, S.H., Xiao, X., and Watkins, S.C. (2006). Alpha-sarcoglycan is recycled from the plasma membrane in the absence of sarcoglycan complex assembly. Traffic 7, 793–810.

26. Li, C., Wilborn, J., Pittman, S., Daw, J., Alonso-Pérez, J., Díaz-Manera, J., Weihl, C.C., and Haller, G. (2023). Comprehensive functional characterization of SGCB coding variants predicts pathogenicity in limb-girdle muscular dystrophy type R4/2E. J. Clin. Invest. 133,.

27. Roberds, S.L., Anderson, R.D., Ibraghimov-Beskrovnaya, O., and Campbell, K.P. (1993). Primary structure and muscle-specific expression of the 50-kDa dystrophin-associated glycoprotein (adhalin). J. Biol. Chem. 268, 23739–23742.

28. Liu, S., Su, T., Xia, X., and Zhou, Z.H. (2025). Native DGC structure rationalizes muscular dystrophy-causing mutations. Nature 637, 1261–1271.

29. Wan, L., Ge, X., Xu, Q., Huang, G., Yang, T., Campbell, K.P., Yan, Z., and Wu, J. (2025). Structure and assembly of the dystrophin glycoprotein complex. Nature 637, 1252–1260.

30. Gastaldello, S., D’Angelo, S., Franzoso, S., Fanin, M., Angelini, C., Betto, R., and Sandonà, D. (2008). Inhibition of proteasome activity promotes the correct localization of disease-causing alpha-sarcoglycan mutants in HEK-293 cells constitutively expressing beta-, gamma-, and delta-sarcoglycan. Am. J. Pathol. 173, 170–181.

31. Carotti, M., Marsolier, J., Soardi, M., Bianchini, E., Gomiero, C., Fecchio, C., Henriques, S.F., Betto, R., Sacchetto, R., Richard, I., et al. (2018). Repairing folding-defective á-sarcoglycan mutants by CFTR correctors, a potential therapy for limb-girdle muscular dystrophy 2D. Hum. Mol. Genet. 27, 969–984.

32. Scano, M., Benetollo, A., Dalla Barba, F., and Sandonà, D. (2024). Advanced therapeutic approaches in sarcoglycanopathies. Curr. Opin. Pharmacol. 76, 102459.

33. Faure, A.J., Schmiedel, J.M., Baeza-Centurion, P., and Lehner, B. (2020). DiMSum: an error model and pipeline for analyzing deep mutational scanning data and diagnosing common experimental pathologies. Genome Biol. 21, 207.

34. Thompson, T.G., Chan, Y.M., Hack, A.A., Brosius, M., Rajala, M., Lidov, H.G., McNally, E.M., Watkins, S., and Kunkel, L.M. (2000). Filamin 2 (FLN2): A muscle-specific sarcoglycan interacting protein. J. Cell Biol. 148, 115–126.

35. Guyon, J.R., Kudryashova, E., Potts, A., Dalkilic, I., Brosius, M.A., Thompson, T.G., Beckmann, J.S., Kunkel, L.M., and Spencer, M.J. (2003). Calpain 3 cleaves filamin C and regulates its ability to interact with gamma- and delta-sarcoglycans. Muscle Nerve 28, 472–483.

36. Yoshida, T., Pan, Y., Hanada, H., Iwata, Y., and Shigekawa, M. (1998). Bidirectional signaling between sarcoglycans and the integrin adhesion system in cultured L6 myocytes. J. Biol. Chem. 273, 1583–1590.

37. Barton, E.R. (2006). Impact of sarcoglycan complex on mechanical signal transduction in murine skeletal muscle. Am J Physiol, Cell Physiol 290, C411–9.

38. Barton, E.R. (2010). Restoration of gamma-sarcoglycan localization and mechanical signal transduction are independent in murine skeletal muscle. J. Biol. Chem. 285, 17263–17270.

39. Jaganathan, K., Kyriazopoulou Panagiotopoulou, S., McRae, J.F., Darbandi, S.F., Knowles, D., Li, Y.I., Kosmicki, J.A., Arbelaez, J., Cui, W., Schwartz, G.B., et al. (2019). Predicting Splicing from Primary Sequence with Deep Learning. Cell 176, 535–548.e24.

40. Cheng, J., Novati, G., Pan, J., Bycroft, C., Þemgulytë, A., Applebaum, T., Pritzel, A., Wong, L.H., Zielinski, M., Sargeant, T., et al. (2023). Accurate proteome-wide missense variant effect prediction with AlphaMissense. Science 381, eadg7492.

41. Danzi, M.C., Dohrn, M.F., Fazal, S., Beijer, D., Rebelo, A.P., Cintra, V., and Züchner, S. (2023). Deep structured learning for variant prioritization in Mendelian diseases. Nat. Commun. 14, 4167.

42. Ioannidis, N.M., Rothstein, J.H., Pejaver, V., Middha, S., McDonnell, S.K., Baheti, S., Musolf, A., Li, Q., Holzinger, E., Karyadi, D., et al. (2016). REVEL: an ensemble method for predicting the pathogenicity of rare missense variants. Am. J. Hum. Genet. 99, 877–885.

43. Dong, C., Wei, P., Jian, X., Gibbs, R., Boerwinkle, E., Wang, K., and Liu, X. (2015). Comparison and integration of deleteriousness prediction methods for nonsynonymous SNVs in whole exome sequencing studies. Hum. Mol. Genet. 24, 2125–2137.

44. Dinh, T., Jang, S.-K., Zaitlen, N., and Ntranos, V. (2026). Compressing the collective knowledge of ESM into a single protein language model. Nat. Methods 23, 772–784.

45. Quinodoz, M., Peter, V.G., Cisarova, K., Royer-Bertrand, B., Stenson, P.D., Cooper, D.N., Unger, S., Superti-Furga, A., and Rivolta, C. (2022). Analysis of missense variants in the human genome reveals widespread gene-specific clustering and improves prediction of pathogenicity. Am. J. Hum. Genet. 109, 457–470.

46. Gao, H., Hamp, T., Ede, J., Schraiber, J.G., McRae, J., Singer-Berk, M., Yang, Y., Dietrich, A.S.D., Fiziev, P.P., Kuderna, L.F.K., et al. (2023). The landscape of tolerated genetic variation in humans and primates. Science 380, eabn8153.

47. Kircher, M., Witten, D.M., Jain, P., O’Roak, B.J., Cooper, G.M., and Shendure, J. (2014). A general framework for estimating the relative pathogenicity of human genetic variants. Nat. Genet. 46, 310–315.

48. Frazer, J., Notin, P., Dias, M., Gomez, A., Min, J.K., Brock, K., Gal, Y., and Marks, D.S. (2021). Disease variant prediction with deep generative models of evolutionary data. Nature 599, 91–95.

49. Qi, H., Zhang, H., Zhao, Y., Chen, C., Long, J.J., Chung, W.K., Guan, Y., and Shen, Y. (2021). MVP predicts the pathogenicity of missense variants by deep learning. Nat. Commun. 12, 510.

50. Jumper, J., Evans, R., Pritzel, A., Green, T., Figurnov, M., Ronneberger, O., Tunyasuvunakool, K., Bates, R., Žídek, A., Potapenko, A., et al. (2021). Highly accurate protein structure prediction with AlphaFold. Nature 596, 583–589.

51. Varadi, M., Anyango, S., Deshpande, M., Nair, S., Natassia, C., Yordanova, G., Yuan, D., Stroe, O., Wood, G., Laydon, A., et al. (2022). AlphaFold Protein Structure Database: massively expanding the structural coverage of protein-sequence space with high-accuracy models. Nucleic Acids Res. 50, D439–D444.

52. Richards, S., Aziz, N., Bale, S., Bick, D., Das, S., Gastier-Foster, J., Grody, W.W., Hegde, M., Lyon, E., Spector, E., et al. (2015). Standards and guidelines for the interpretation of sequence variants: a joint consensus recommendation of the American College of Medical Genetics and Genomics and the Association for Molecular Pathology. Genet. Med. 17, 405–424.

53. Brnich, S.E., Abou Tayoun, A.N., Couch, F.J., Cutting, G.R., Greenblatt, M.S., Heinen, C.D., Kanavy, D.M., Luo, X., McNulty, S.M., Starita, L.M., et al. (2019). Recommendations for application of the functional evidence PS3/BS3 criterion using the ACMG/AMP sequence variant interpretation framework. Genome Med. 12, 3.

54. Alonso-Pérez, J., González-Quereda, L., Bello, L., Guglieri, M., Straub, V., Gallano, P., Semplicini, C., Pegoraro, E., Zangaro, V., Nascimento, A., et al. (2020). New genotype-phenotype correlations in a large European cohort of patients with sarcoglycanopathy. Brain 143, 2696–2708.

55. Luce, L., Kocak, G.S., Verdú-Díaz, J., Alonso-Pérez, J., Claeys, K.G., Stojkovic, T., Fernández-Eulate, G., Laforêt, P., Miladi, N., Di Pace, F., et al. (2026). Cracking the Code: Genotype-Phenotype Correlation Models in Sarcoglycanopathies. Ann. Clin. Transl. Neurol.

56. Xie, Z., Hou, Y., Yu, M., Liu, Y., Fan, Y., Zhang, W., Wang, Z., Xiong, H., and Yuan, Y. (2019). Clinical and genetic spectrum of sarcoglycanopathies in a large cohort of Chinese patients. Orphanet J. Rare Dis. 14, 43.

57. (2023). Targeted massive parallel sequencing and comprehensive phenome-genome assessment in neuromuscular patients without genetic etiology by conventional diagnostic methods - 조안나 - 서울대학교 : 논문 - DBpia.

58. Shimazaki, R., Saito, Y., Awaya, T., Minami, N., Kurosawa, R., Hosokawa, M., Ohara, H., Hayashi, S., Takeuchi, A., Hagiwara, M., et al. (2025). Profiling of pathogenic variants in Japanese patients with sarcoglycanopathy. Orphanet J. Rare Dis. 20, 1.

59. Riccio, C., Jansen, M.L., Guo, L., and Ziegler, A. (2024). Variant effect predictors: a systematic review and practical guide. Hum. Genet. 143, 625–634.

60. Karchin, R., Radivojac, P., O’Donnell-Luria, A., Greenblatt, M.S., Tolstorukov, M.Y., and Sonkin, D. (2024). Improving transparency of computational tools for variant effect prediction. Nat. Genet. 56, 1324–1326.

61. Hord, J.M., Turk, R., Kusano, H., Rader, E.P., Burns, S., Gastel, Z., Prouty, S.J., Yu, L., Burden, S.J., and Campbell, K.P. (2026). Cytoplasmic region of beta-dystroglycan is essential for postsynaptic maturation and neuromuscular function in mice. Proc Natl Acad Sci USA 123, e2600931123.

62. Nagy, E., and Maquat, L.E. (1998). A rule for termination-codon position within intron-containing genes: when nonsense affects RNA abundance. Trends Biochem. Sci. 23, 198–199.

63. Wang, J., Gudikote, J.P., Olivas, O.R., and Wilkinson, M.F. (2002). Boundary-independent polar nonsense-mediated decay. EMBO Rep. 3, 274–279.

64. Maquat, L.E. (2004). Nonsense-mediated mRNA decay: splicing, translation and mRNP dynamics. Nat. Rev. Mol. Cell Biol. 5, 89–99.

65. Hug, N., Longman, D., and Cáceres, J.F. (2016). Mechanism and regulation of the nonsense-mediated decay pathway. Nucleic Acids Res. 44, 1483–1495.

66. 66. Jagannathan, S., Shadle, S., Resnick, R., Snider, L., Tawil, R.N., van der Maarel, S.M., Bradley, R.K., and Tapscott, S.J. (2016). Model systems of DUX4 expression recapitulate the transcriptional profile of FSHD cells. Hum. Mol. Genet.

67. Gauthier, L.O., Wang, Z., Ng, K.K., Huang, S., Mao, Y., Lek, M., and Ma, K. (2025). Protocol to implement saturation mutagenesis-reinforced functional assays to resolve small-sized variants in disease-related genes. STAR Protocols 6, 103909.

