## Supplemental Figures for "Comprehensive Mapping of *SGCA* Variant Effects Reveals Domain-Specific Constraints Relevant to Sarcoglycanopathies"

Figure S1

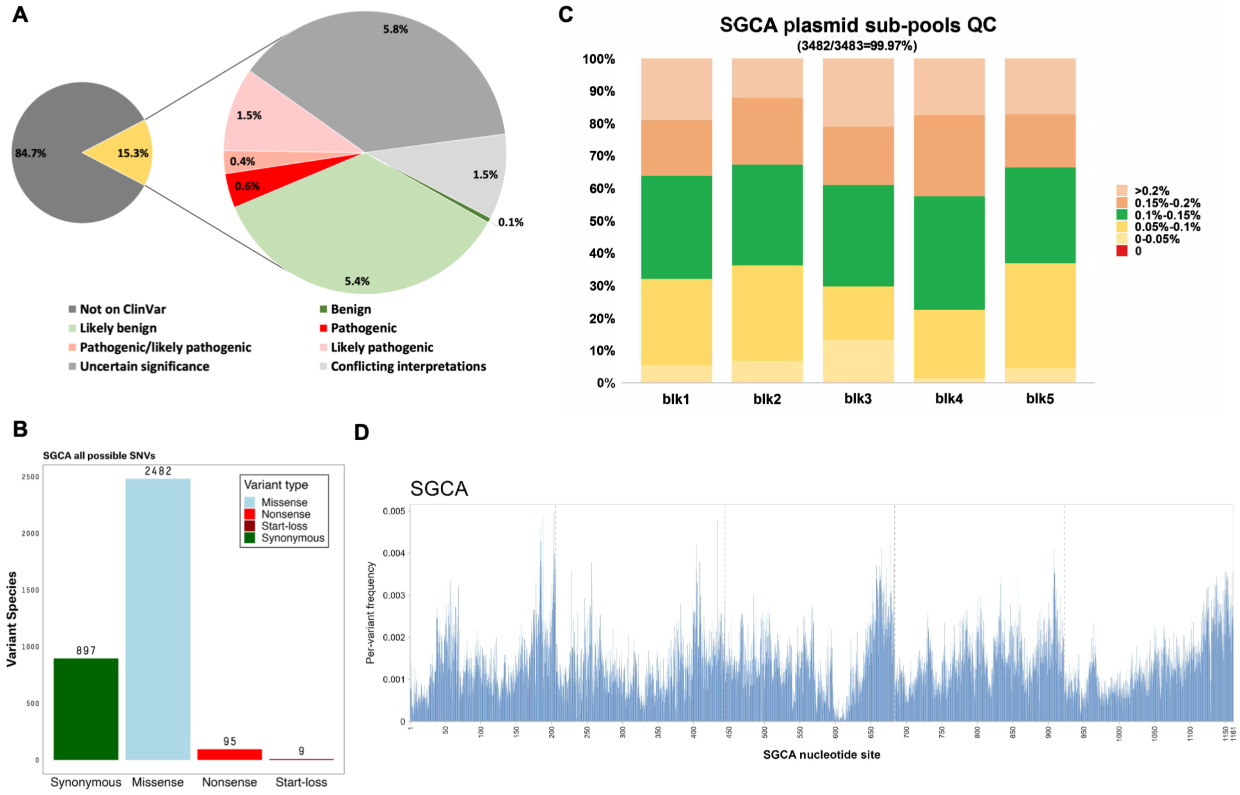


**Figure S1. Classification of coding single nucleotide variants (SNVs) in the *SGCA* gene and quality assessment of the saturation mutagenesis library.**

**(A)** Distribution of *SGCA* coding SNVs in ClinVar. Clinical variant annotations were retrieved from ClinVar (downloaded June 14th, 2026). The left pie chart shows the proportion of all possible SNVs (3,483) represented in ClinVar (533), and the expanded pie chart shows their clinical classifications.

**(B)** Composition of all possible *SGCA* SNVs by variant consequence based on the *SGCA* CDS (NM_000023.4).

**(C)** Variant representation in the five *SGCA* plasmid sub-pools assessed by Amplicon-EZ sequencing. 3,482 of 3,483 possible coding SNVs (99.97%) were detected. Colors indicate the relative representation of individual variants within each sub-pool. The single variant (c.496G>A) not detected in this initial library QC was subsequently detected in all three biological replicates of the functional assay, although at lower coverage, and was therefore assigned a low-confidence score.

**(D)** Per-variant frequency distribution across the *SGCA* CDS in the plasmid sub-pools. The x-axis represents nucleotide positions along the *SGCA* CDS, with dashed vertical lines indicating the boundaries between the five mutagenesis blocks.

Figure S2

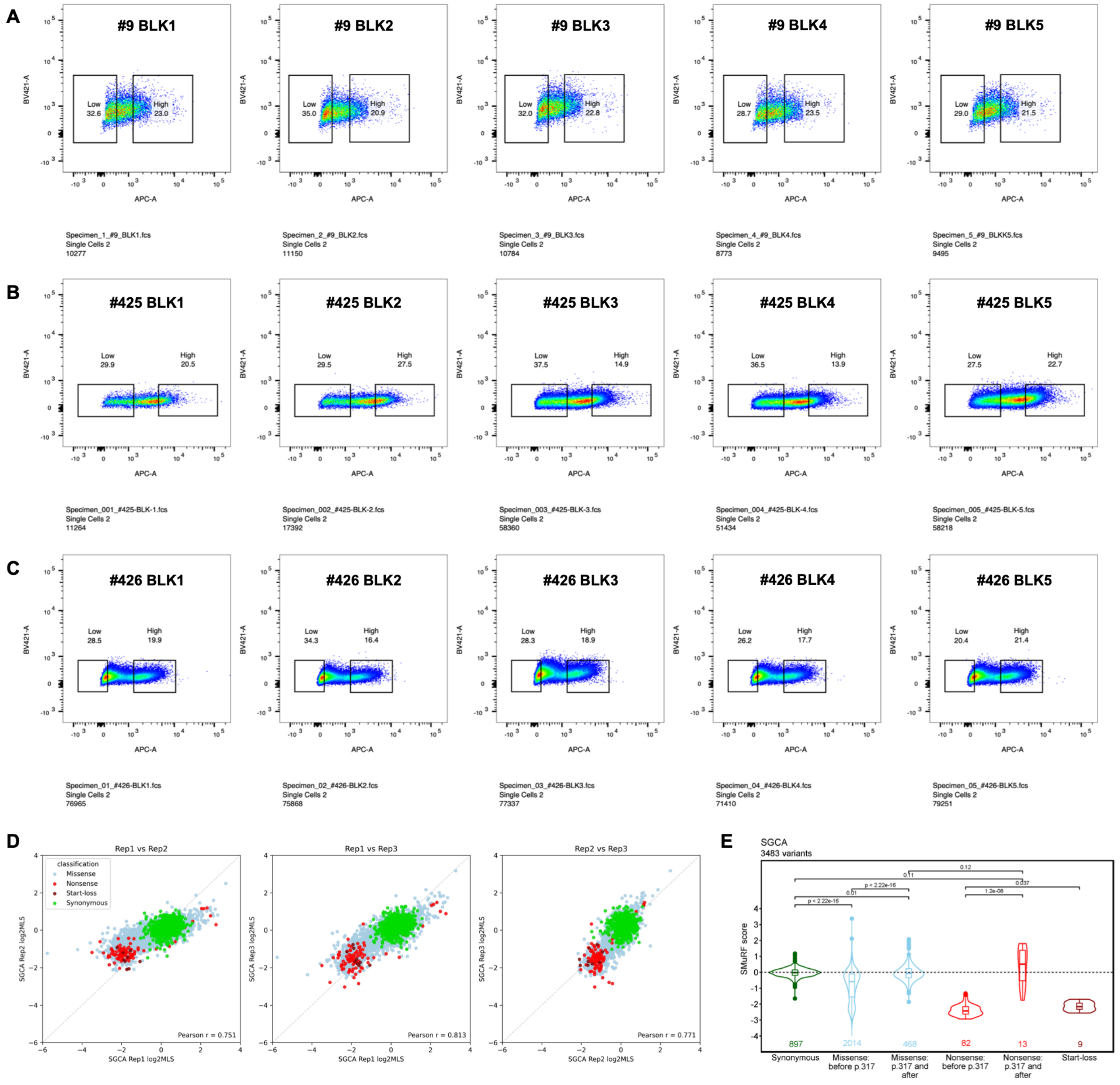


**Figure S2. Quality control and reproducibility of the *SGCA* SMuRF assay.**

**(A-C)** Representative fluorescence-activated cell sorting (FACS) plots showing gating of high- and low-surface-expression populations based on FLAG-APC fluorescence intensity in HEK-BDG #9 (A), HEK-BDG #425 (B), and HEK-BDG #426 (C) stable cell lines across all five mutagenesis blocks (BLK1-BLK5). High- and low-surface-expression populations were selected to enrich for approximately the top 20% and bottom 30% of FLAG-positive cells, respectively. Gating boundaries were adjusted slightly between experiments to accommodate differences in fluorescence distributions while maintaining comparable population sizes. For each block, at least 20,000 events were recorded to determine gating parameters. Cell debris, dead cells, and multiplets were excluded prior to sorting.

**(D)** Reproducibility of SGCA membrane localization measurements across biological replicates. Pairwise correlations of replicate-specific log2 membrane localization scores (log2MLS) for high-confidence *SGCA* variants (n = 3,398) derived from three independent biological replicates. Each point represents a single variant and is colored by variant class. Pearson correlation coefficients are shown for each pairwise comparison. The dashed gray line denotes the line of identity (y = x).

**(E)** Distribution of SMuRF scores for all 3,483 possible *SGCA* coding single-nucleotide variants (SNVs), grouped by variant consequence and protein position. Missense and nonsense variants are further stratified by their location before residue 317 or at and after residue 317. The box boundaries represent the 25th and 75th percentiles, with the horizontal line indicating the median and whiskers extending to 1.5 × the interquartile range (IQR). Variant counts are shown below each violin. P values were calculated using two-sided Wilcoxon tests.

Figure S3

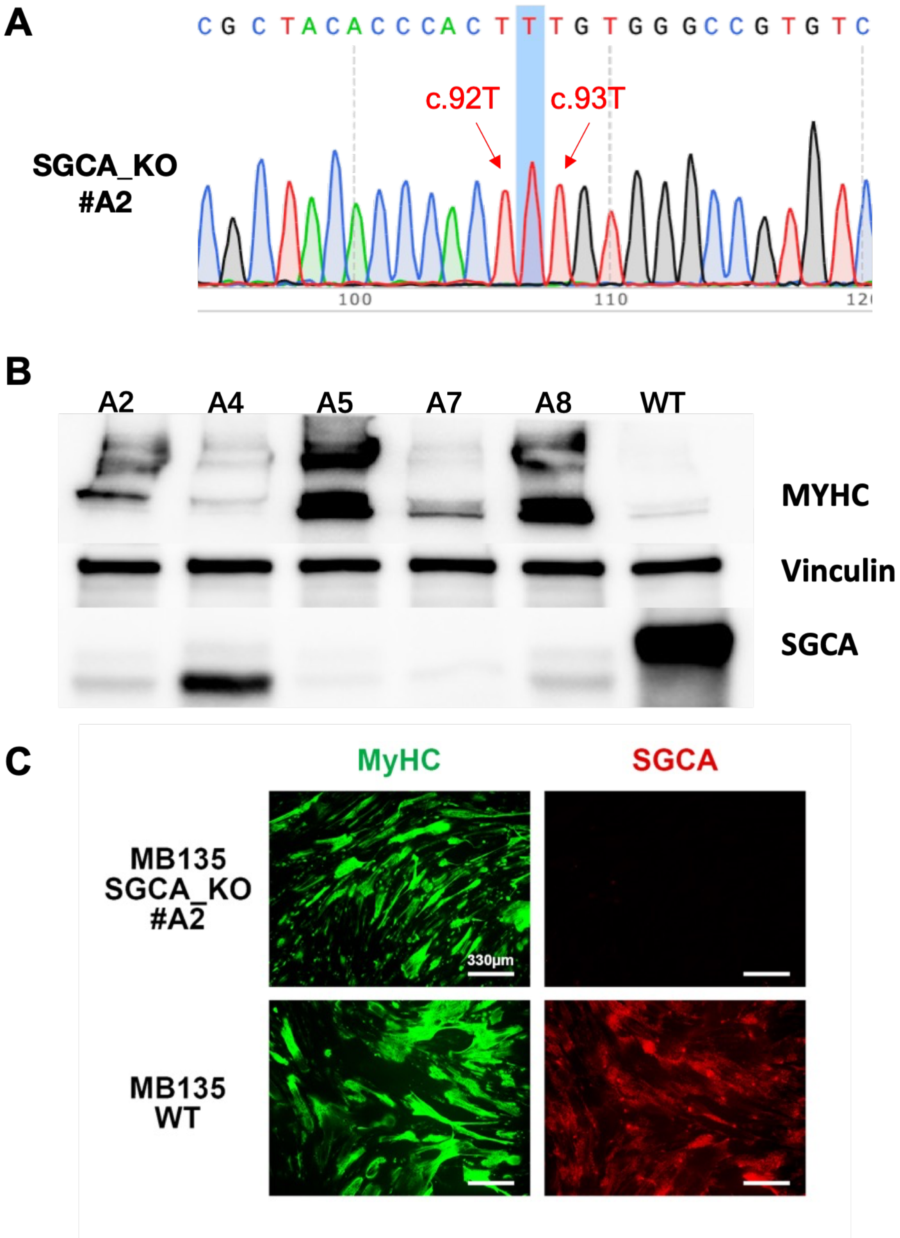


**Figure S3. Validation of *SGCA*-knockout MB135 myoblasts.**

**(A)** Representative Sanger sequencing chromatogram confirming the homozygous c.92dupT variant in the MB135 *SGCA*-knockout (KO) clone A2 generated by CRISPR-Cas9 genome editing.

**(B)** Western blot validation of *SGCA* knockout in MB135 myoblast-derived myotubes. Cell lysates from wild-type MB135 cells and independent *SGCA*-KO clones were analyzed by immunoblotting for SGCA. MyHC was used to confirm myogenic differentiation and vinculin served as the loading control. Anti-SGCA (Invitrogen, MA5-37982, 1:1,000), anti-

MyHC (Invitrogen, 14-6503-37, 1:500), and anti-vinculin (Sigma-Aldrich, V9131, 1:80,000) were used as primary antibodies.

**(C)** Immunofluorescence analysis of MB135 wild-type and *SGCA*-KO (clone A2) myotubes. Cells were differentiated into myotubes, fixed, and stained for MyHC (green) and SGCA (red). Anti-SGCA (Invitrogen, MA5-37982, 1:400) and anti-MyHC (Invitrogen, 14-6503-37, 1:500) were used as primary antibodies. Nuclei were counterstained with DAPI. Scale bars, 330 µm.

Figure S4

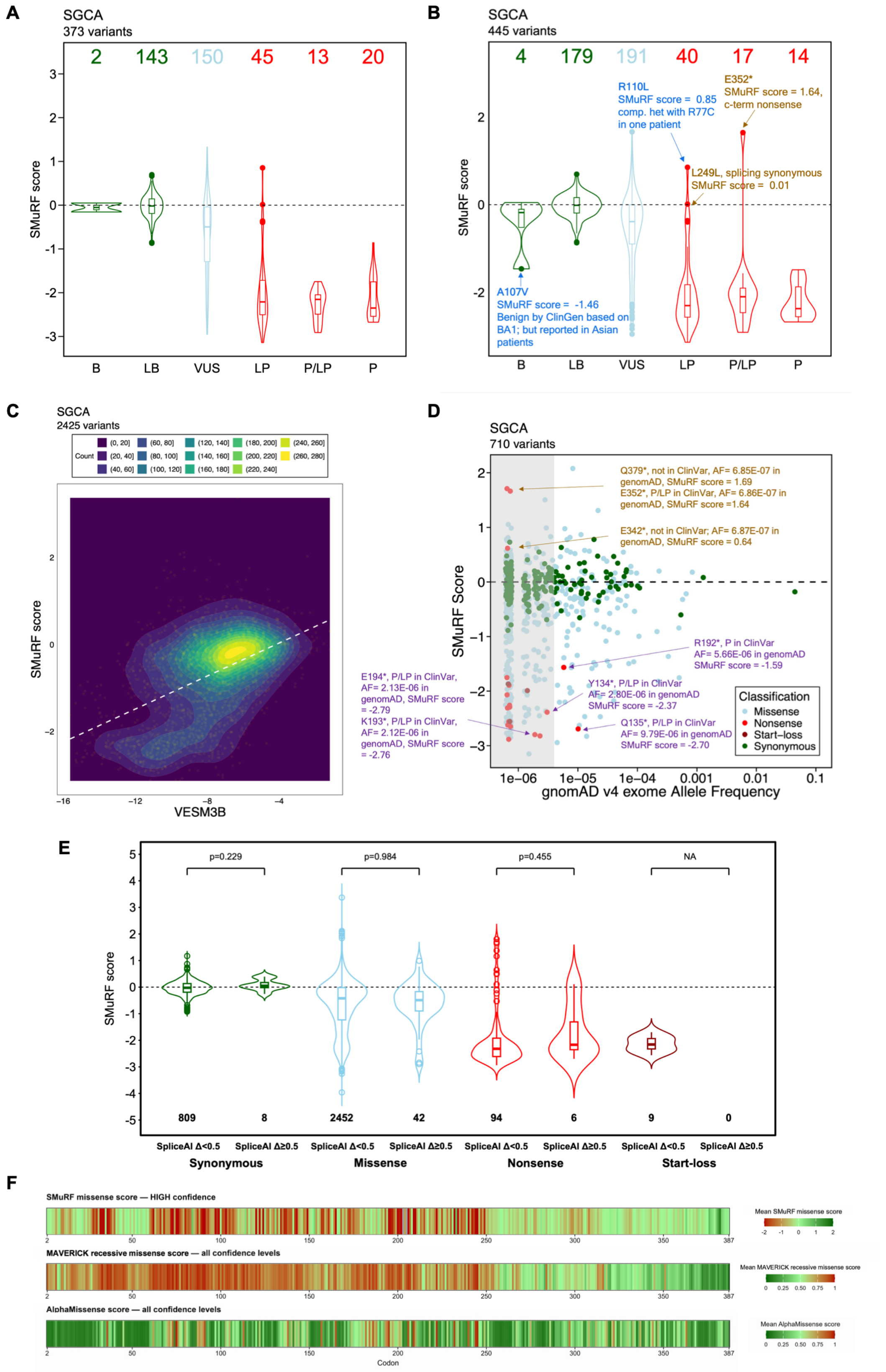

A
SGCA
373 variants
2 143 150 45 13 20
SMuRF score
B LB VUS LP P/LP P
B
SGCA
445 variants
4 179 191 40 17 14
R110L
SMuRF score = 0.85
comp. het with R77C
in one patient
E352*
SMuRF score = 1.64,
c-term nonsense
L249L, splicing synonymous
SMuRF score = 0.01
A107V
SMuRF score = -1.46
Benign by ClinGen based on
BA1; but reported in Asian
patients
C
SGCA
2425 variants
Count
VESM3B
D
SGCA
710 variants
Q379*, not in ClinVar, AF= 6.85E-07 in genomAD, SMuRF score = 1.69
E352*, P/LP in ClinVar, AF= 6.86E-07 in genomAD, SMuRF score =1.64
E342*, not in ClinVar; AF= 6.87E-07 in genomAD, SMuRF score = 0.64
R192*, P in ClinVar
AF= 5.66E-06 in genomAD
SMuRF score = -1.59
Y134*, P/LP in ClinVar
AF= 2.80E-06 in genomAD
SMuRF score = -2.37
Q135*, P/LP in ClinVar
AF= 9.79E-06 in genomAD
SMuRF score = -2.70
E194*, P/LP in ClinVar, AF= 2.13E-06 in genomAD, SMuRF score = -2.79
K193*, P/LP in ClinVar, AF= 2.12E-06 in genomAD, SMuRF score = -2.76
Classification
Missense
Nonsense
Start-loss
Synonymous
SMuRF Score
gnomAD v4 exome Allele Frequency
E
p=0.229 p=0.984 p=0.455 NA
SMuRF score
809 8 2452 42 94 6 9 0
SpliceAI Δ<0.5 SpliceAI Δ≥0.5
Synonymous Missense Nonsense Start-loss
F
SMuRF missense score — HIGH confidence
Mean SMuRF missense score
MAVERICK recessive missense score — all confidence levels
Mean MAVERICK recessive missense score
AlphaMissense score — all confidence levels
Mean AlphaMissense score
Codon

**Figure S4. Additional analyses supporting the clinical relevance of *SGCA* SMuRF scores.**

**(A)** SMuRF scores grouped by ClinVar classification after excluding variants at and beyond residue 317. Variant counts are shown above each violin. The box boundaries represent the 25th and 75th percentiles, with the horizontal line indicating the median and whiskers extending to 1.5 × the interquartile range (IQR). Excluding cytoplasmic-domain variants resulted in stronger concordance between SMuRF scores and ClinVar classifications. Clinical variant annotations were obtained from the ClinVar release downloaded on June 14, 2026.

**(B)** ClinVar-classified variants with discordant SMuRF scores. Representative outlier variants discussed in the text are annotated, including A107V, R110L, L249L, and E352*. Clinical variant annotations from the ClinVar release downloaded on February 2, 2026 were used in this panel to retain A107V as a benign variant prior to its subsequent reclassification. The remaining highlighted outliers are unchanged from those shown in Figure 3A.

**(C)** Density plot showing the relationship between VESM3B scores and SMuRF scores for high-confidence *SGCA* missense variants. Color intensity represents local variant density. White dashed lines represent linear regression.

**(D)** Relationship between SMuRF scores and gnomAD v4 exomes allele frequencies. Representative outlier variants are annotated. The gray shaded region indicates variants with AF < 4e-6. Variants are colored according to variant class. The dashed line indicates the wild-type SMuRF score. Points were jittered for visualization.

**(E)** Comparison of SMuRF scores between variants with low (SpliceAI $\Delta < 0.5$) and high (SpliceAI $\Delta \geq 0.5$) predicted splice impact, stratified by variant consequences. Violin plots show the distribution of SMuRF scores, with box boundaries representing the 25th and 75th percentiles, the horizontal line indicating the median, and whiskers extending to 1.5 × the interquartile range (IQR). Variant counts are shown below each violin. P values were calculated using two-sided Wilcoxon tests.

**(F)** Comparison of residue-level mean SMuRF missense scores, mean MAVERICK recessive missense scores, and mean AlphaMissense scores across the *SGCA* coding sequence. For each codon, the mean score of all high-confidence SMuRF missense substitutions or all possible computationally predicted missense substitutions was displayed as a heatmap. Color intensity represents the residue-level mean missense scores for each method.

Figure S5

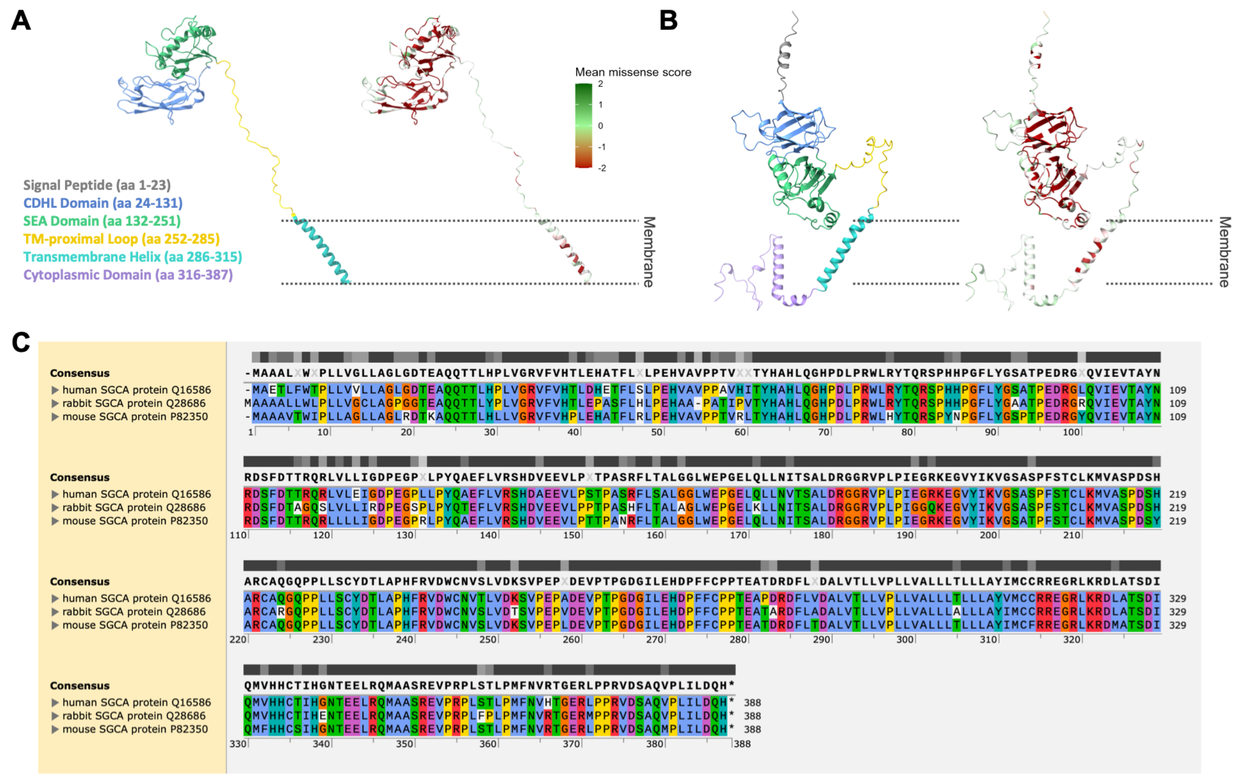


**Figure S5. Comparison of experimentally determined and predicted SGCA structures and sequence conservation across species.**

**(A)** Mouse SGCA cryo-EM structure colored by structural domains (left) or mean missense SMuRF score (right). Similar to the rabbit SGCA structure used in Figure 4A, the experimentally resolved mouse structure includes the extracellular region and transmembrane helix but does not fully resolve the cytoplasmic domain.

**(B)** AlphaFold-predicted human SGCA structure colored by structural domains (left) or mean missense SMuRF score (right). The predicted model provides a structural representation of the complete protein, including the signal peptide and cytoplasmic domain that are absent or incompletely resolved in the experimentally determined structures.

**(C)** Multiple sequence alignment of human (UniProt Q16586), rabbit (UniProt Q28686), and mouse (UniProt P82350) SGCA protein sequences generated using Clustal Omega and visualized in SnapGene. Amino acids are colored according to physicochemical properties, and sequence conservation is indicated by the grayscale bars above the alignment.

Figure S6

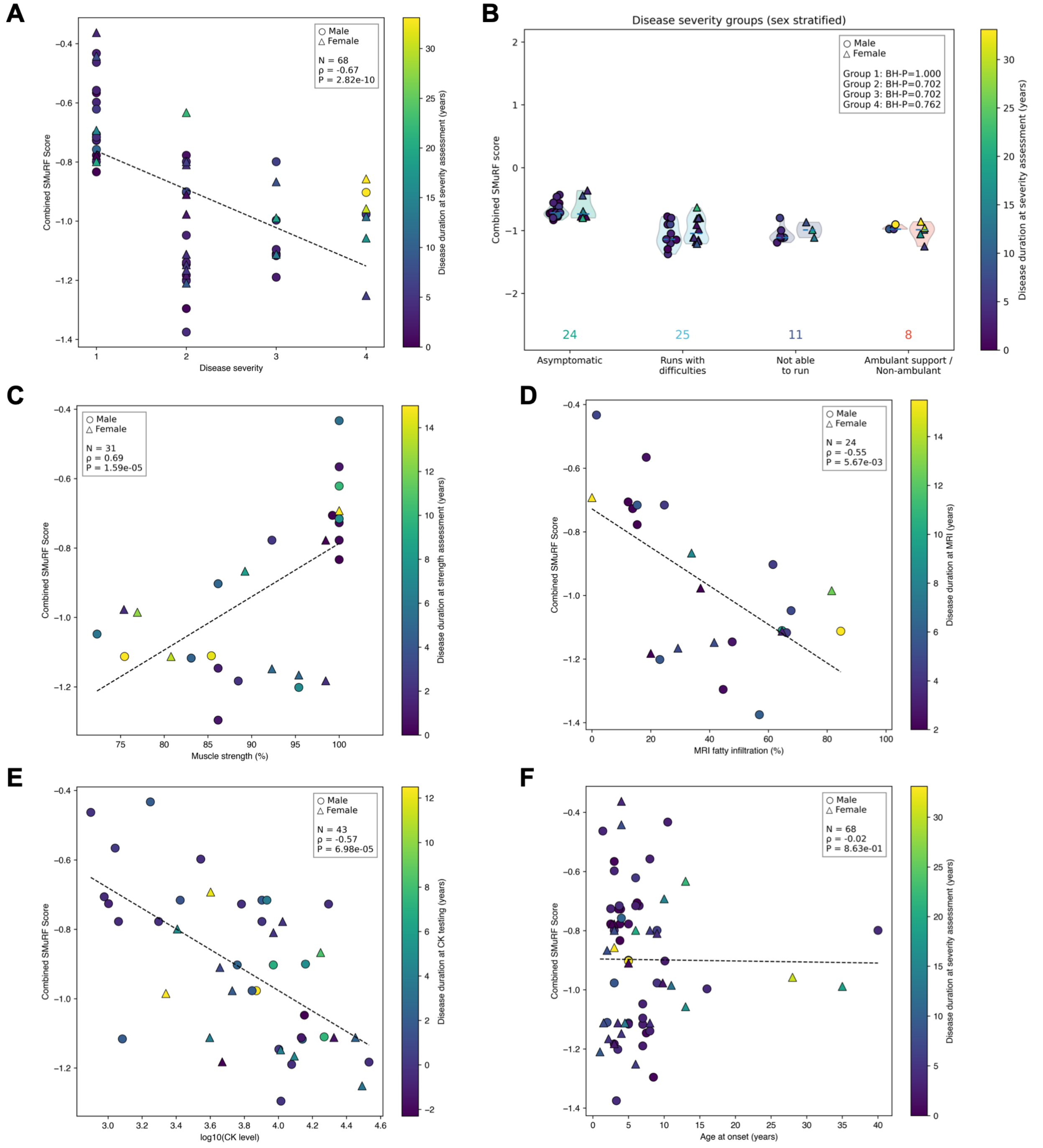


**Figure S6. Correlation of combined SMuRF scores with clinical measures of disease severity.**

**(A)** Correlation between combined SMuRF scores and clinical disease severity. Each point represents one patient and is colored according to disease duration at the time of severity assessment.

**(B)** Combined SMuRF scores stratified by clinical disease severity and sex. Patients were categorized as: (1) asymptomatic hyperCKemia or minimally affected, (2) ambulatory with difficulty running, (3) ambulatory but unable to run, and (4) ambulatory with support or non-ambulant. Within each severity group, combined SMuRF scores were compared between males and females. Violin plots show score distributions, and dashed blue lines indicate medians. P values for sex comparisons were adjusted using the Benjamini-Hochberg method. Numbers below each group indicate the total sample size.

**(C-F)** Correlation between combined SMuRF scores and muscle strength (C), MRI fatty infiltration (D), serum creatine kinase (CK) levels (E), and age at disease onset (F), respectively. CK levels are shown as log10-transformed values.

Individual symbols represent participants (circles, males; triangles, females), with symbol colors indicating disease duration at the time of the corresponding clinical assessment. Spearman's rank correlation coefficients (ρ) and two-sided P values are shown in (A) and (C-F). In (B), P values for male-versus-female comparisons within each disease severity group were calculated using two-sided Mann-Whitney U tests and adjusted for multiple comparisons using the Benjamini-Hochberg procedure.
